# Genes controlling polysaccharide synthesis, protein folding and basic metabolism ensure intra-host cell adaptation and replication of *Yersinia enterocolitica*

**DOI:** 10.64898/2026.09.16.752068

**Authors:** Jiline Brinkmann, Jiabin Huang, Patrick Blümke, Martin Aepfelbacher, Klaus Ruckdeschel

**Affiliations:** Institute for Medical Microbiology, Virology and Hygiene, University Medical Center Eppendorf; Leibniz Institute of Virology, Hamburg Martinistr. 52, 20251 Hamburg, Germany

## Abstract

Pathogenesis of yersiniosis comprises cell-invasive events in which the bacteria subvert the endocytic-autophagolysosomal pathway of infected host cells to create an intracellular niche for bacterial replication. To determine potential virulence factors that enable intracellular proliferation of *Yersinia enterocolitica*, we performed a high-throughput transposon-directed insertion site sequencing approach (TraDIS) with infected epithelial HeLa cells. In this screen, we identified seven particular genes that have either metabolic activities in disulfide bond formation (*dsbA*), glutamine synthesis (*glnA*), and energy generation (*acnB*), or dictate synthesis of enterobacterial common antigen ECA (*webB/C/F*) and LPS (*wzx*). Individual knockout of the identified genes impaired intracellular replication of *Yersinia*. Furthermore, these mutants differentially affected host cell autophagy and the acidification of *Yersinia*-containing vacuoles (YCVs). The results indicate that *glnA*, *acnB*, *dsbA*, *wecB*, and *wecC* support autophagy-promoted, intracellular replication of Yersinia. Folding of yet to be defined proteins dependent on *dsbA* seems to be most critically required for mediating these events. A *wzx*-controlled LPS synthesis pathway appears implicated in preventing YCV acidification. These results suggest that autophagy induction and adjustment of *Yersinia* to the intravacuolar environment result from a multifactorial adaptation process that involves formation of a protective carbohydrate envelope and basic metabolic enzymes needed for the refinement of productive intracellular replication.

## INTRODUCTION

Three species of the genus *Yersinia* are pathogenic for humans and rodents. These are *Y. pestis*, the causative agent of plague, and *Y. enterocolitica* and *Y. pseudotuberculosis* which mediate gastrointestinal syndromes, such as enteritis and mesenteric lymphadenitis. These *Yersinia* species share common virulence mechanisms conferred by the plasmid-encoded Ysc type III protein secretion system which is highly conserved in *Y. pestis*, *Y. pseudotuberculosis* and *Y. enterocolitica* ^1, 2^. The Ysc type III secretion system (T3SS) mediates the translocation of Yop effector proteins into host cells where the Yops modify cellular immune responses ^1, 2^. This enables extracellular survival of the bacteria in the host lymphoid tissues. *Yersinia* is consequently generally considered as extracellular pathogen.

However, the infectious lifestyle of the pathogenic yersiniae also involves cell invasion and replication in the interior of infected cells^1–4^. This seems important especially in the initial stages of infection^3, 4^. Accordingly, the enteropathogenic species *Y. enterocolitica* and *Y. pseudotuberculosis* engage the outer membrane protein invasin to gain access to the intestinal mucosa after ingestion by contaminated food or water^5, 6^. Invasin binds and activates β1-integrin receptors exposed on the surface of eukaryotic cells^7, 8^. This triggers bacterial internalization and uptake of yersiniae into *Yersinia*-containing vacuoles (YCVs), in which yersiniae are able to survive. Consequently, β1-integrin-mediated uptake by intestinal cells supports transmigration of *Yersinia* through the intestinal mucosa to reach the submucosal host lymphoid tissue of the Peyers patches^3, 6, 8^. Likewise, the pathogenesis of plague involves survival within host cells, particularly in macrophages^4, 9^. This suggests that not only extracellular survival in the lymphoid tissue mediated by the Ysc secretion system, but also intracellular uptake and replication in host cells are common features in the strategies of pathogenic *Yersinia* species to establish infection and mediate disease. *Yersinia* species may therefore be seen as facultative intracellular pathogens^4^.

The mechanisms that allow intracellular survival and replication of yersiniae within host cells are, however, not well understood. It has been shown that yersiniae modulate endosome recycling to create the protective YCV in which the bacteria can multiply^4, 9^. The YCV membrane rapidly acquires the late endosomal / lysosomal marker Lamp1, but subsequent maturation to a degrative lysosomal compartment is impaired in a substantial subset of YCVs^10, 11^. In these YCVs, the bacteria are protected against acidification and proteolysis, which enables their intracellular replication. Importantly, several studies have shown that the formation of such proliferative YCVs involves execution / activation of autophagic processes in host cells^11–14^. Accordingly, these YCVs display autophagy-related features, such as LC3 processing, LC3 recruitment, and double membrane formation. Our previous studies have shown that fusion of these autophagic YCVs with lysosomes is actively prevented by *Y. enterocolitica* in epithelial cells^11^. This results in proliferation of Y. enterocolitica in the autophagic compartments. The study indicated that *Y. enterocolitica* exploits host cell autophagy to generate a protective niche for intracellular bacterial replication^11^. Interestingly, autophagy induction and proliferation of *Yersinia* in the autophagic YCV occur independently of the *Yersinia* virulence plasmid and of Ysc-dependent type III secretion^11–14^. *Yersinia* factors encoded on the chromosome may instead play a major role in mediating the intracellular lifestyle of *Yersinia*^4^.

In line, several chromosomal factors had been reported to be implicated in enabling intracellular survival of yersiniae in host cells. It appears that proteins of the general bacterial stress response essentially contribute to the adaption of *Yersinia* to the intracellular environment. Accordingly, the stress-responsive PhoPQ two-component system is critically involved^10, 15^. PhoP transcriptionally regulates *mgtC* and *ugd*, which are supposed to provide protection against low Mg++ conditions and antimicrobial peptides, respectively^10^. Furthermore, the osmotic regulator OmpR and the periplasmic protease GsrA play role in adjusting yersiniae to changes in osmolarity and to other not yet defined stressful conditions in the YCV^16, 17^. The superoxide dismutase SodA and the manganese transporter MntH are required to protect the bacteria from intravacuolar oxidative stress^18, 19^. A transposon site hybridization screen had furthermore identified genes encoding for enzymes of the enterobacterial common antigen ECA (*wecB*, *wecC*) and the lipooligosaccharide (*galU*) biosynthesis pathway to be important for intracellular survival of *Y. pestis*^20^. ECA and lipooligosaccaride in the bacterial envelope likely increase the resistance to antimicrobial peptides and to other potentially toxic substances in the YCV^20^. Similarly, the multifaceted membrane protein OmpA may aid to confer protection against antimicrobial peptides^21^. From these findings it appears that intracellular proliferation of *Yersinia* in host cells is a multifactorial process that depends on an array of bacterial stress response elements and on structural components of the bacterial envelope. The understanding of the genetic basis for intracellular proliferation of Yersinia appears nevertheless still incomplete^4^.

In order to learn more about the *Yersinia* factors and mechanisms implicated in intracellular multiplication, we applied a transposon-directed insertion site sequencing TraDIS approach on epithelial HeLa cells infected with *Y. enterocolitica*. High throughput transposon insertion sites sequencing of bacterial pools before and after cell infection identified several conditionally essential genes involved in intracellular replication. Analysis of individual knockout mutants of these genes indicates that there is an overall coherence between autophagy induction and increased intracellular bacterial replication. It was verified that ECA of *Yersinia* plays role in enabling efficient intracellular survival. Besides ECA also an intact LPS structure seems to be required for adequate intracellular multiplication as a *Yersinia* strain mutated in the O-antigen subunit flippase Wzx is severely impaired in intracellular survival. Furthermore, metabolic activities depending on the glutamine synthetase GlnA and the aconitase AcnB contribute to intracellular *Yersinia* proliferation. In addition, the thiol-disulfide oxidoreductase DsbA, which mediates proper folding of bacterial envelope and secreted proteins, is critically involved. The obtained results support the concept that multiple *Yersinia* factors from different levels of activities concertedly avert the intravacuolar antibacterial responses for efficient bacterial replication.

## RESULTS

### Transposon-directed insertion site sequencing (TraDIS) discloses seven genes with critical importance for intracellular survival of *Y. enterocolitica*

Y. *enterocolitica* induces the formation of YCVs with autophagic characteristics and prevents YCV fusion with lysosomes which protects the enclosed bacterial population from degradation^11, 13, 14^. The detailed mechanisms and specific virulence factors influencing these processes are, however, largely unclear. This study therefore aimed to establish a *Y. enterocolitica* transposon mutant library and use it for infection and high-throughput sequencing to identify bacterial genes that play role in ensuring the intracellular survival of *Yersinia*. As, according to previous studies, the intracellular lifestyle of *Yersinia* mainly depends on chromosomal factors and does not require the Yersinia pYV virulence plasmid and Ysc type III protein secretion^4^, we used a *Yersinia* strain that lacked the pYV plasmid for transposon mutagenesis. The strain was electroporated with a transposon-transposase complex and mutants were selected by kanamycin resistance. The obtained colonies were pooled to create the mutant library, which finally contained around 100 000 clones. The entire mutant library was used for infection of epithelial HeLa cells and a large-scale intracellular survival assay was performed (Fig. 1A). Viable, surviving mutants were then subjected to TraDIS and compared to the control samples. In total, four different pools of mutants were analysed by TraDIS. Sample 1 was the initial input library obtained after pooling all the transformed clones (Fig. 1A, “initial library”). Essential genes that cannot be mutated without causing death of the respective clone should have been lost in sample 1. In sample 2, the initial input library was sub-cultured in bacterial growth medium for two hours to enrich vital mutants (“pre-infection library”). This sample allows comparison of the mutant populations at the beginning and after infection. The pre-grown pre-infection pool of mutants was then used for HeLa cell infection. In parallel, a portion of the pre-infection mutant library was grown for 24 hours in cell culture medium in the absence of cells to eliminate mutants with general growth and fitness deficits that do not result from challenges within the host cell (sample 3, “outgrowth control library”). The infection of HeLa cells was conducted at a relatively low MOI of 20 to limit bottleneck effects and direct competition between the mutants. Gentamicin was added to kill extracellular bacteria that had not invaded the cells after one hour of infection^22^. After a final incubation time of 24 hours, the cells were lysed to release and enrich surviving intracellular mutants (sample 4 in Fig. 1A: “intracellular mutant library”).

**Fig. 1:**
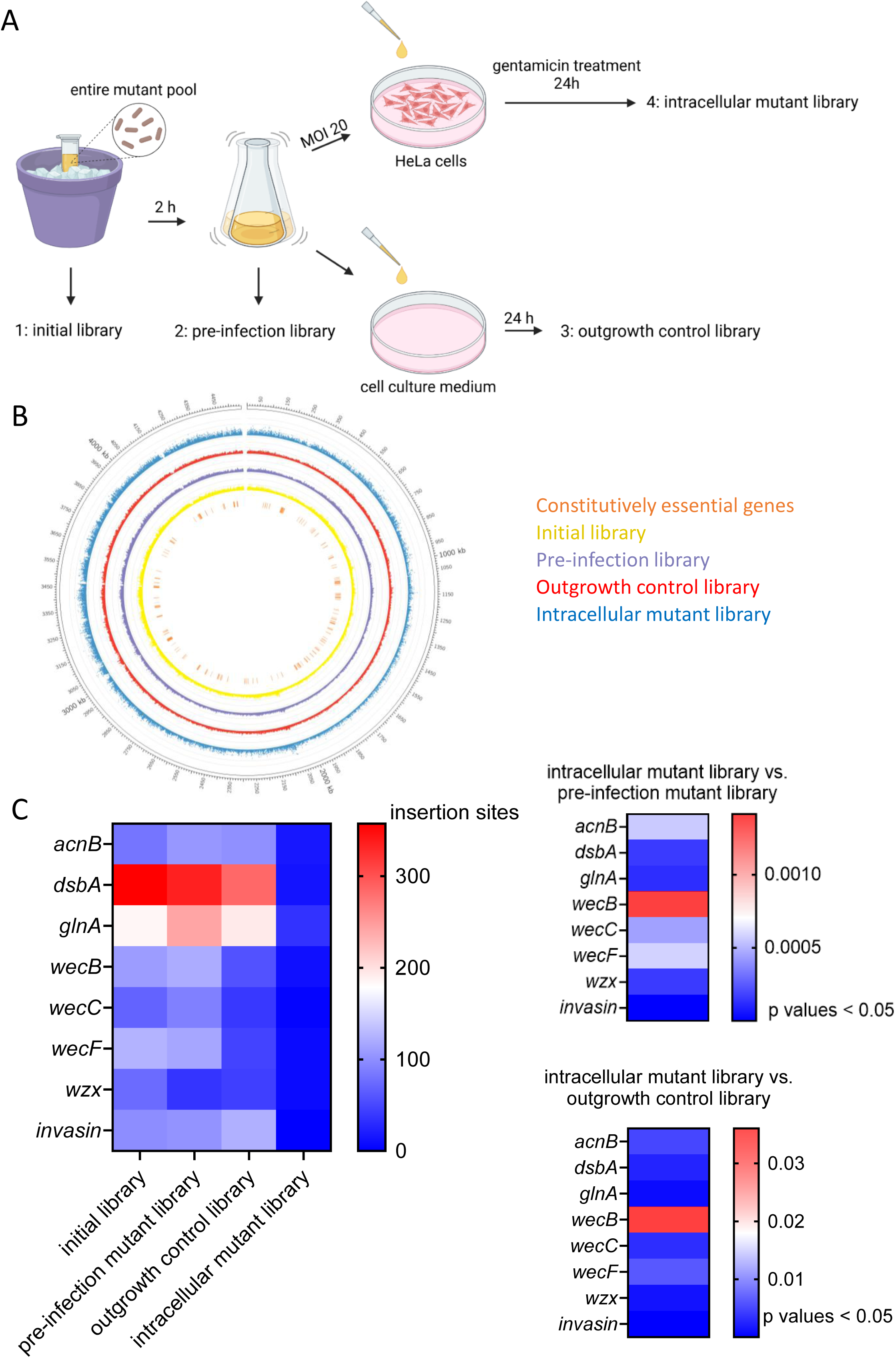
Set-up and results of the high-throughput TraDIS experiment. A. Procedure of sample preparation: The first sample (1) encompassed all transposon mutants directly obtained after transformation and kanamycin resistance selection (initial library). The second sample (2) was the pre-infection mutant pool after subculturing in bacterial growth medium for 2 h (pre-infection library). The third sample (3) contained mutants that grew in cell culture medium for 24 h (outgrowth control library). The fourth sample (4) were the viable intracellular bacteria released from HeLa cells infected with Yersinia for 1 h and then incubated in presence of gentamicin for all in all 23 h (intracellular mutant library). B. Ideogram of the chromosome of *Y. enterocolitica* WA-314 with identified transposon insertion sites: Each circle represents the normalized template counts for each transposon insertion site and experimental condition. Each dot marks one sequenced unique templates. The colours correspond to the following: yellow = initial library (1), purple = pre-infection library (2), red = outgrowth control library (3), blue = intracellular mutant library (4). The identified essential genes (∼217) are labelled in orange. The scale marks kilobases within the genome. C. Heatmap of insertion sites per sample: Identified conditional essential genes were significantly less frequently mutated in the sample containing intracellular bacteria than in the outgrowth control and the pre-infection mutant library. The mean numbers of insertion sites per gene is displayed by colour (left panel). The adjusted p-values of the genes that significantly differed between the intracellular mutant library and the pre-infection library (upper right panel) and outgrowth control library (lower right panel) are displayed by colour.

To identify all transposon insertion sites in each mutant genome, high-throughput sequencing was performed. Paired-end short-read sequencing identified the genetic region adjacent to the transposon. The obtained sequences were aligned with the *Yersinia enterocolitica subsp. enterocolitica* WA-314 genome (GenBank accession number CP009367) by bioinformatic analysis. More than 20 million transposon-containing reads were sequenced per sample (Supplemental table 1). The number of insertion sites was determined for each open reading frame (ORF). A total of 120,738 insertion sites were found distributed across the entire genome (Fig. 1B). Since the *Y. enterocolitica* genome consists of approximately 4.6 million base pairs and encodes about 4,000 genes, 30-fold coverage was achieved, with an average mutation site occurring every 38 bp.

The constitutive essentiality of genes for bacterial viability was assessed by identifying genes with significant absence of transposon insertions in the initial mutant library (sample 1). Approximately 217 genes were determined to be constitutively essential, representing about 5.4% of the entire genome (Supplemental table 2). Typical essential genes required for bacterial viability were ribosomal proteins (*rpmJ*, *rpsM*, *rpsK*, *rpsD*, *rplQ*, *rpmI*, *rplT*), translation factors (*infC*, *pheS*/*pheT*), as well as proteins involved in DNA replication (*dnaA*, *dnaN*), cell division (*ftsH*), protein translocation (*secY*, *secD*, *secF*), nucleoid organization (*ihfA*, *ihfB*), ribosome maturation (*rimP*, *rbfA*), central metabolism (*glmM*), and tRNA processing (*thrS*). These genes are universally essential across bacterial species^23^, which supports the validity of the experimental setup and the analysis pipeline.

To determine genes that may play a conditional role in the intracellular lifestyle of *Yersinia*, we compared the data of the intracellular mutant library (sample 4) with the pre-infection library (sample 2) and the outgrowth control library (sample 3). ORFs that were significantly less frequently mutated in the intracellular mutant library, with an adjusted p-value of less than 0.05, were considered conditional genes required for the intracellular survival of *Y. enterocolitica* in epithelial host cells. The genes that differed significantly from both the pre-infection and the outgrowth control library were *dsbA*, *glnA*, *inv*, *acnB*, *wecB*, *wecC*, *wecF*, and *wzx* (Fig. 1C, Supplemental table 3). It is important to note that *invasin* (*inv*) was among the identified genes. Invasin confers host cell adherence and bacterial internalization, which is prerequisite for intracellular survival of *Yersinia*^6, 7^. The detection of *invasin* verifies the applied experimental procedure as appropriate approach to identify conditional genes indispensable for intracellular survival of *Yersinia*. The other identified conditional genes (Table 1) could be assigned to two main groups. First, there are two enzymes predominantly involved in metabolic processes, which are glutamine synthetase A (GlnA) and aconitase B (AcnB). Then there are proteins related to the synthesis of outer membrane components. The Wec proteins and the CH47_2457 gene product, a Wzx flippase (Genbank accession No. AAC60766), play role in the synthesis and export of ECA and LPS, respectively. The disulfide bond interchange protein A DsbA may also belong to the second group. DsbA is located in the periplasm and ensures proper folding of multiple membrane and exterior protein by disulfide bond formation^24^.

**Table 1:** List of genes that identified in the TraDIS screen to be conditionally essential for the intracellular survival of *Y. enterocolitica* with their predicted molecular functions.

| Name | Molecular function |
| --- | --- |
| *glnA* | glutamine synthetase type I |
| *acnB* | bifunctional aconitate hydratase 2/2-methylisocitrate dehydratase |
| *wecB* | UDP-N-acetylglucosamine 2-epimerase, synthesis of enterobacterial common antigen |
| *wecC* | UDP-N-acetyl-D-mannosamine dehydrogenase, synthesis of enterobacterial common antigen |
| *wecF* | 4-alpha-L-fucosyltransferase, synthesis of enterobacterial common antigen |
| *wzx* | flippase, polysaccharide biosynthesis family protein, membrane protein involved in the export of O-antigen |
| *dsbA* | disulfide interchange protein, periplasmic protein disulfide isomerase I |

### Targeted knockout of the identified genes impairs intracellular replication of *Yersinia*

To determine whether the conditional genes identified in the TraDIS screen may indeed play role in intracellular survival and proliferation of *Yersinia*, we specifically knocked out the respective genes by deleting the entire genomic sequence via CRISPR-assisted homology repair. The obtained mutants were first characterized in their general growth behaviour in bacterial and cell culture growth medium. The mutants did not exhibit significant growth defects in both media, with the exception of the Δ*wecB* mutant, which displayed noticeable reduced growth in cell culture medium (Suppl. Fig. 1A).

Then, the intracellular survival of the mutants was investigated. For this purpose, a gentamicin protection assay with HeLa cells was performed related to the screening workflow shown in Fig. 1A. The cells were infected for 30 min, fresh medium with gentamicin was then added to kill extracellular bacteria, and cells were lysed at 1 and 24 h post infection (p. i.) to determine the number of intracellular bacteria. All mutants had comparable sensitivities towards gentamicin as assessed by disc diffusion assay (Suppl. Fig. 1B). The replication rate was calculated by dividing the number of bacteria at 24 h p.i. by the initial number of bacteria at 1 h p. i.. The 1 h values were used to determine the initial rates of cell infection by the different strains. All mutants were capable of interacting with HeLa cells and did not significantly differ in the numbers of cell-associated bacteria after 1 h (Suppl. Fig. 2A). Similarly, the rates of cell internalization were comparable for the investigated Yersinia strains, as determined by double immunofluorescence staining of bacteria before and after cell permeabilization (Suppl. Fig. 2B, 2C).

The *Y.a enterocolitica* WT strain WAc (lacking the pYV plasmid) replicated well inside cells; its replication rate was 8.9 at the 24 h time point (Fig. 2A). An *E. coli* strain that expresses the *Yersinia* invasin protein (*E. coli* +inv) was used as negative control^25^. This strain can enter host cells via the same uptake mechanism as *Yersinia*. However, it is unable to trigger autophagy and is eliminated by Hela cells^11^. Accordingly, the replication rate of *E. coli* +inv was 0.1. Importantly, also a mutant lacking DsbA did not survive inside the cells. Its number of CFUs decreased from 1 h p. i. to 24 h p. i. and the corresponding replication rate was 0.1 (Δ*dsbA*). The other mutants could survive in HeLa cells, but were impaired in intracellular replication. As such, they multiplied less efficiently than the WAc WT strain, with proliferation rates of 1.9 for the Δ*glnA* mutant, 2.2 for the Δ*acnB* mutant, 2.0 for the Δ*wecB* mutant, 4.1 for the Δ*wecC* mutant, 6.9 for the Δ*wecF* mutant, and 1.1 for the Δ*wzx* mutant. With the exception of the Δ*wecF* mutant, these differences to WAc were statistically significant. Thus, the data obtained with the individual knockout mutants largely validate the results of TraDIS screening approach, indicating that the identified genes play a critical role in mediating the intracellular survival and replication of *Yersinia*.

**Fig. 2:**
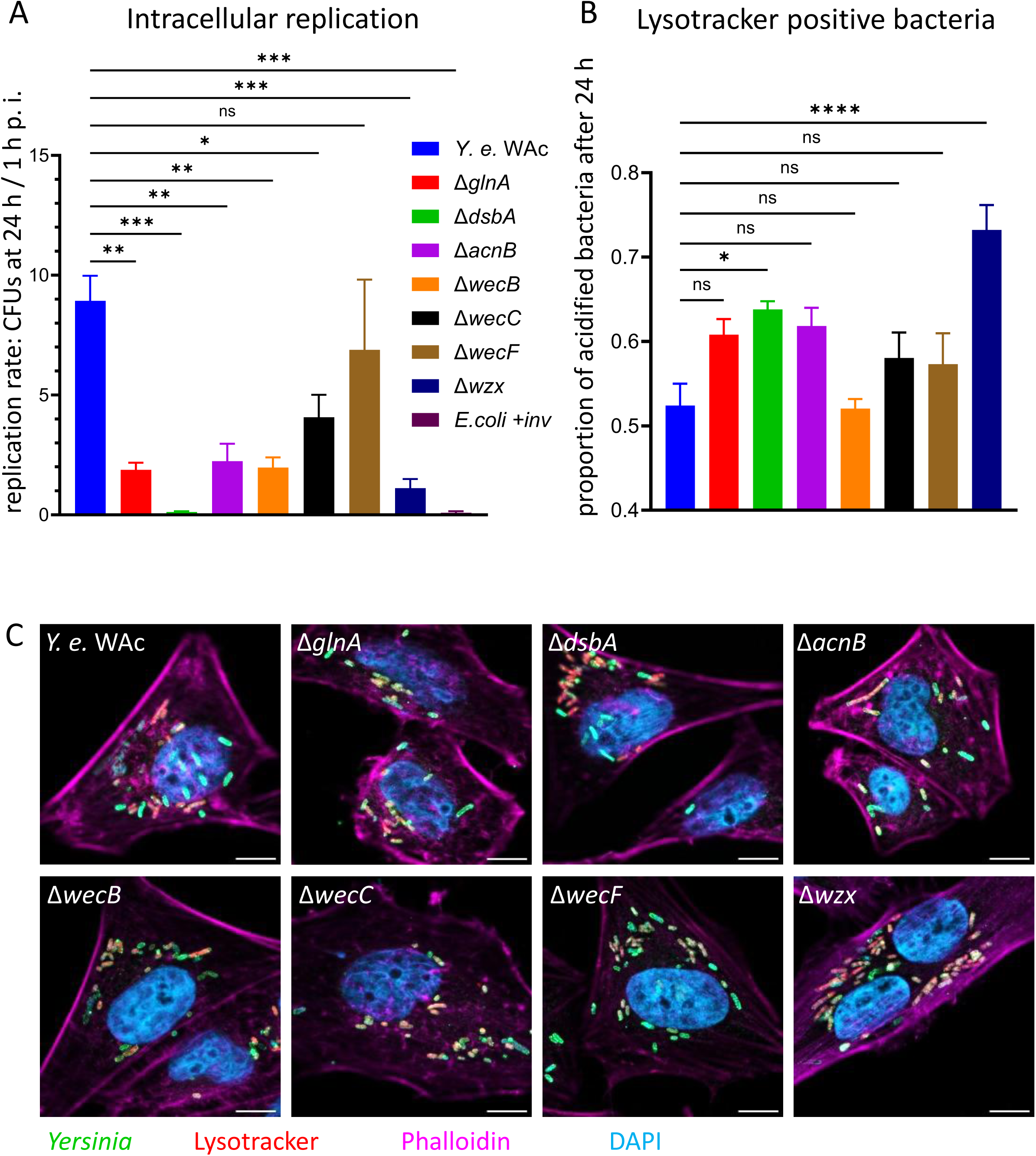
Intracellular bacterial replication and YCV acidification after *Yersinia* infection. A. Analysis of intracellular survival and proliferation of the constructed mutants by gentamicin protection assay: HeLa cells were infected for 30 min, then treated with gentamicin, and CFUs of intracellularly surviving bacteria were determined after 1 and 24 h p. i. The replication rates were calculated by dividing the numbers of CFUs at 24 h p. i. by the numbers of CFUs at 1 h p. i.. Three individual experiments were performed and evaluated. B. Quantification of Lysotracker Red-positive (acidified) bacteria inside HeLa cells at 24 h p. i.: The areas of all bacteria and Lysotracker-positive bacteria inside the cells were examined. The ratios were calculated by dividing the Lysotracker-positive bacteria by the total population of bacteria. Three individual experiments were performed and five images of each were quantified.. Representative microscopy images of Lysotracker staining: HeLa cells were infected and stained with Lysotracker Red 30 min prior fixation at 24 h p. i.. Immunostaining was performed to label the bacteria with an LPS antibody in green. Phalloidin stained actin in far-red and DAPI was used to mark bacterial DNA and nuclear DNA of the infected cells. Coloured images of all channels are merged. Scale bars denote 10 μm.

To learn more about the intracellular fate of the internalized yersiniae we used the dye Lysotracker to stain acidic compartments. The acquisition of Lysotracker positivity is indicative for subjection of the respective YCV to the degradative lysosomal pathway. In accordance with our previous studies (ref Maria) we found that about half of the intracellular *Yersinia* WAc WT population (52.4 %) was acidified 24 h p. i. (Fig. 2B, 2C). The WAc mutants that were impaired in intracellular replication, but yet survived in HeLa cells (Δ*acnB*, Δ*wecB*, Δ*wecC*, Δ*glnA*, Δ*wecF*) exhibited only modestly higher percentages of acidified bacteria. But notably, the strains Δ*dsbA* and Δ*wzx*, which cannot survive inside the cells (Fig. 2A), produced significantly higher proportions of Lysotracker-positive bacteria (63.81 % for Δ*dsbA*, and 73.21 % for Δ*wzx*). The enhanced intracellular acidification of Δ*dsbA* and Δ*wzx* in HeLa cells consequently suggests that these mutants are likely eliminated through the lysosomal pathway.

### DsbA, GlnA, AcnB, WecB and WecC contribute to the formation of protective autophagic vacuoles

Our previous studies have shown that the formation of proliferative YCVs depends on the acquisition of autophagy-related characteristics^11^. We therefore investigated the capabilities of the *Yersinia* mutants to induce autophagy in HeLa cells.

One way to monitor autophagy is analysing the conversion of LC3 I to LC3 II by immunoblotting^26^. LC3 I is processed to LC3 II when incorporated in the autophagosomal membrane. HeLa cells were infected with *Yersinia* WAc WT, the knockout mutants, or *E. coli* +*inv*, and the levels of cytosolic LC3B I and lipidated LC3B II were determined in cellular lysates 4 h p. i.. The intensities of the LC3B-I and LC3B-II bands were measured and the LC3 conversion ratios were calculated by dividing the LC3B-II values by those for LC3B-I. Finally, the obtained results were normalized to the *Yersinia* WT WAc reference strain to determine the autophagy induction rate in relation to WAc.

The knockout mutants Δ*glnA*, Δ*dsbA*, Δ*acnB*, Δ*wecB*, and Δ*wecC* induced significantly less LC3 conversion than the WAc WT (Fig. 3A, 3B). Accordingly, they converted LC3B at approximately 50% of the rate of the wild type, which is similar to the value of the *E. coli* +*inv* strain. The data on LC3 conversion correlated well with the recruitment of LC3 to the YCVs. This was assessed in HeLa cells that stably express a LC3 version tagged with both RFP and GFP. The RFP-GFP-LC3 fusion construct allows visualisation of the maturation of autophagosomes (neutral pH, yellow colour) to autolysosomes (acidic pH, red colour) because the GFP signal is lost upon lysosomal fusion and resultant acidification^27^. The percentages of bacteria in RFP-GFP-LC3-positive vacuoles were microscopically determined after 4 h of infection. Approximately 20% of intracellular WAc WT bacteria localized into RFP-GFP-LC3-positive vacuoles (Fig. 3C, 3D). The mutants Δ*glnA*, Δ*dsbA*, Δ*acnB*, Δ*wecB*, and Δ*wecC* were found to be significantly less frequently LC3-positive compared to WAc. This went along with reduced LC3 processing (Fig. 3B). The strain Δ*wecF*, however, did not significantly differ from WAc in mediating LC3 conversion (Fig. 3B) and LC3 recruitment (Fig. 3C). This was in concordance with only slightly reduced intracellular replication of Δ*wecF* in comparison to WAc (Fig. 2A). These results confirm a notable correlation between the capacities of these strains to induce autophagy and to multiply intracellularly, which supports the concept that the activation of autophagic events is critically involved in enabling the intracellular lifestyle of *Yersinia*^11^. The bacteria in the RFP-GFP-LC3-positive vacuoles exhibited for the most part simultaneous GFP and RFP fluorescence 4 h p. i., which is reflected by green to yellow colour in Fig. 3D. This supports the evidence that survival of *Yersinia* in autophagic compartments is associated with blockage of autophagosome to autolysosome maturation^11^. Together, these data suggest that GlnA, DsbA, AcnB, WecB and WecC contribute to the formation of protective autophagic YCVs and concomitant intracellular survival and proliferation of *Yersinia*. Interestingly, the Δ*wzx* mutant differed from this coherence. This strain was not impaired in autophagy induction, as indicated by high rates of LC3 conversion (Fig. 3B) and LC3 decoration (Fig. 3C), but nevertheless was unable to multiply inside the cells (Fig. 2A). The Δ*wzx*-containing YCVs were significantly subjected to acidification at later stages of infection (Fig. 2B), which suggests that blockage of the autophagic flux towards autolysosome formation by the Δ*wzx* mutant may be insufficient, finally leading to Δ*wzx* elimination.

**Fig. 3:**
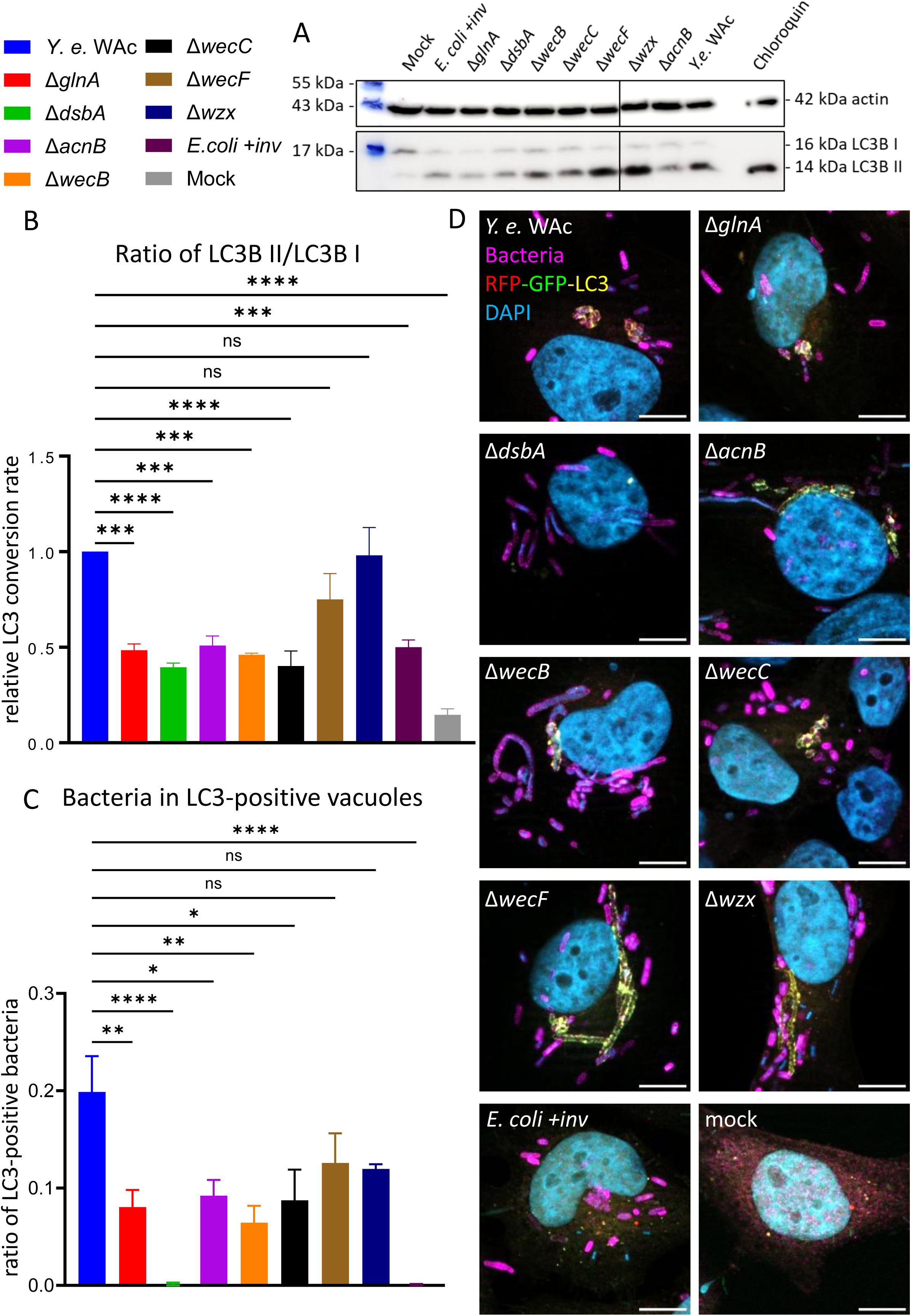
Autophagy induction and formation of LC3 positive YCVs after *Yersinia* infection. A. Conversion of LC3B I to LC3B II determined by Western blotting: HeLa cells were infected as indicated and the levels of LC3B and actin were analysed in cell lysates 4 h p. i. with specific primary antibodies. A representative western blot image is shown. B. Quantification of LC3B I to LC3B II conversion: To monitor and quantify autophagy induction, the band intensities of LC3 I and LC3 II obtained after immunoblotting (as in A) were measured. The LC3 conversion rates were then calculated by dividing the intensity of LC3 II by LC3 I for each investigated condition. Three individual experiments were analysed. C. Quantification of LC3-positive bacteria in HLC3-HeLa cells: HLC3 cells stably expressing RFP-GFP-LC3 were infected and the numbers of DAPI-stained bacteria and bacteria inside LC3-positive vacuoles were counted at 4 h p. i.. The ratios were calculated by dividing the numbers of LC3-positive bacteria by the total bacterial numbers. Three individual experiments were performed and ten images were quantified. D. Representative microscopy images of LC3-positive YCVs: HLC3 cells expressing RFP-GFP-LC3 were infected and fixed at 4 h p. i.. Immunostaining was performed to label the bacteria with a LPS antibody dyed in far-red. DAPI was used to stain bacterial and nuclear DNA of the human cells. Coloured images of all channels are merged. Scale bars denote 10 μm.

### DsbA is essential for autophagy induction and intracellular replication of *Yersinia*

The Δ*dsbA* mutant exhibited the most pronounced phenotype with respect to strongly impaired autophagy and compromised intracellular survival. To verify the importance of DsbA in mediating these events in *Yersinia* infection, we reconstituted the Δ*dsbA* mutant with the homologous *dsbA* sequence in fusion with an alfa-tag motif. The expression of DsbA by the complemented mutant was confirmed by Western blotting (Fig. 4A). Importantly, the reconstitution of the Δ*dsbA* mutant with DsbA largely restored the intracellular survival of the mutant (Fig. 4B). In line, the recruitment of LC3 to the YCVs was restored by complementing the Δ*dsbA* mutant (Fig. 4C). These results support the assumption that DsbA plays a crucial role in ensuring autophagy induction and mediating the intracellular survival of *Yersinia*.

**Fig. 4:**
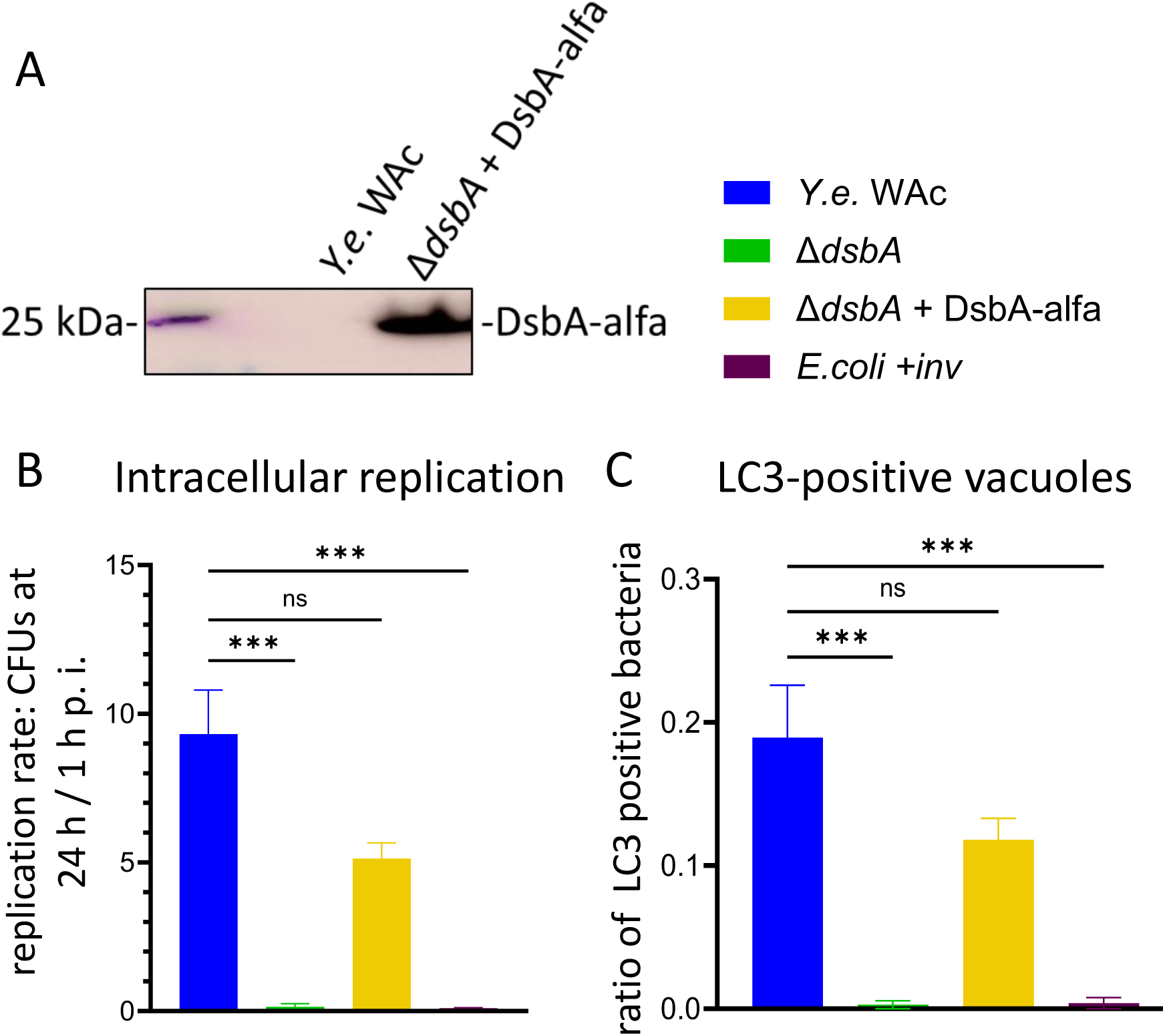
Restoration of intracellular *Yersinia* survival and LC3 recruitment by *dsbA* reconstitution. A. Reconstitution of the Δ*dsbA* mutant with DsbA-alfa: The DsbA-alfa construct in the complemented Δ*dsbA* mutant was verified by western blotting using anti-alfa-tag antibody. The detected protein band corresponds to the expected size of 25 kDa.. Analysis of intracellular replication by gentamicin protection assay: HeLa cells were infected for 30 min with the indicated strains, then treated with gentamicin, and CFUs of intracellularly surviving bacteria were determined after 1 and 24 h p. i.. The replication rates were calculated by dividing the numbers of CFUs at 24 h p. i. by the numbers of CFUs at 1 h p. i.. Three individual experiments were performed and evaluated. C. Quantification of LC3-positive bacteria in HLC3 cells: HLC3-HeLa cells stably expressing RFP-GFP-LC3 cells were infected and the numbers of DAPI-stained bacteria and bacteria inside LC3-positive vacuoles were counted 4 h p. i.. The ratios were calculated by dividing the numbers of LC3-positive bacteria by the total bacterial numbers. Three individual experiments were performed and ten images were quantified. Representative microscopy images are displayed in Fig. 5D.

To characterize the involvement of DsbA in autophagy induction in more detail, we investigated the recruitment of upstream autophagy-initiating signaling molecules to the YCVs. Previous results have shown that autophagy-inducing yersiniae first localize into Lamp1-positive YCVs that are then decorated with ubiquitin and p62^11^. Ubiquitin marks and conveys targets to autophagic degradation through interaction with p62^28, 29^. p62 subsequently contacts LC3 for autophagy initiation.

The RFP-GFP-LC3 expressing HeLa cells were infected with WAc, the Δ*dsbA* mutant, the reconstituted strain Δ*dsbA* +DsbA-alpha, and the *E. coli* +*inv* strain. Additionally, we investigated the Δ*wzx* mutant which triggered autophagy (Fig. 3), but was impaired in intracellular survival (Fig. 2). The cells were infected for 4 h and then stained for endogenous ubiquitin, p62, and Lamp1. The numbers of bacteria positive for LC3, ubiquitin, p62, and Lamp1 were quantified (Fig. 5). As expected, Lamp1 was recruited to the majority of the intracellular bacteria, also to WAc Δ*wzx* and *E. coli* +*inv* bacteria which did not reside in LC3-labeled YCVs (Fig. 5A, 5D). Lamp1 positivity was, however, slightly reduced for the Δ*dsbA* mutant. The accumulation of ubiquitin and p62 on the other hand correlated well with the mobilization of RFP-GFP-LC3 to the intracellular bacteria. Accordingly, the strains that localized into LC3-positive YCVs (WAc, Δ*dsbA* +DsbA-alpha, Δ*wzx*) were also able to recruit ubiquitin and p62 (Fig. 5B, 5C, 5D), with co-localization rates of ubiquitin with RFP-GFP-LC3 from 76 and 82 %, and of p62 with RFP-GFP-LC3 from 98 and 100 %. Thus, autophagy mediated by WAc, the reconstituted mutant Δ*dsbA* +DsbA-alpha, and Δ*wzx* seems to succeed through classical, selective autophagy involving ubiquitin and p62.

**Fig. 5:**
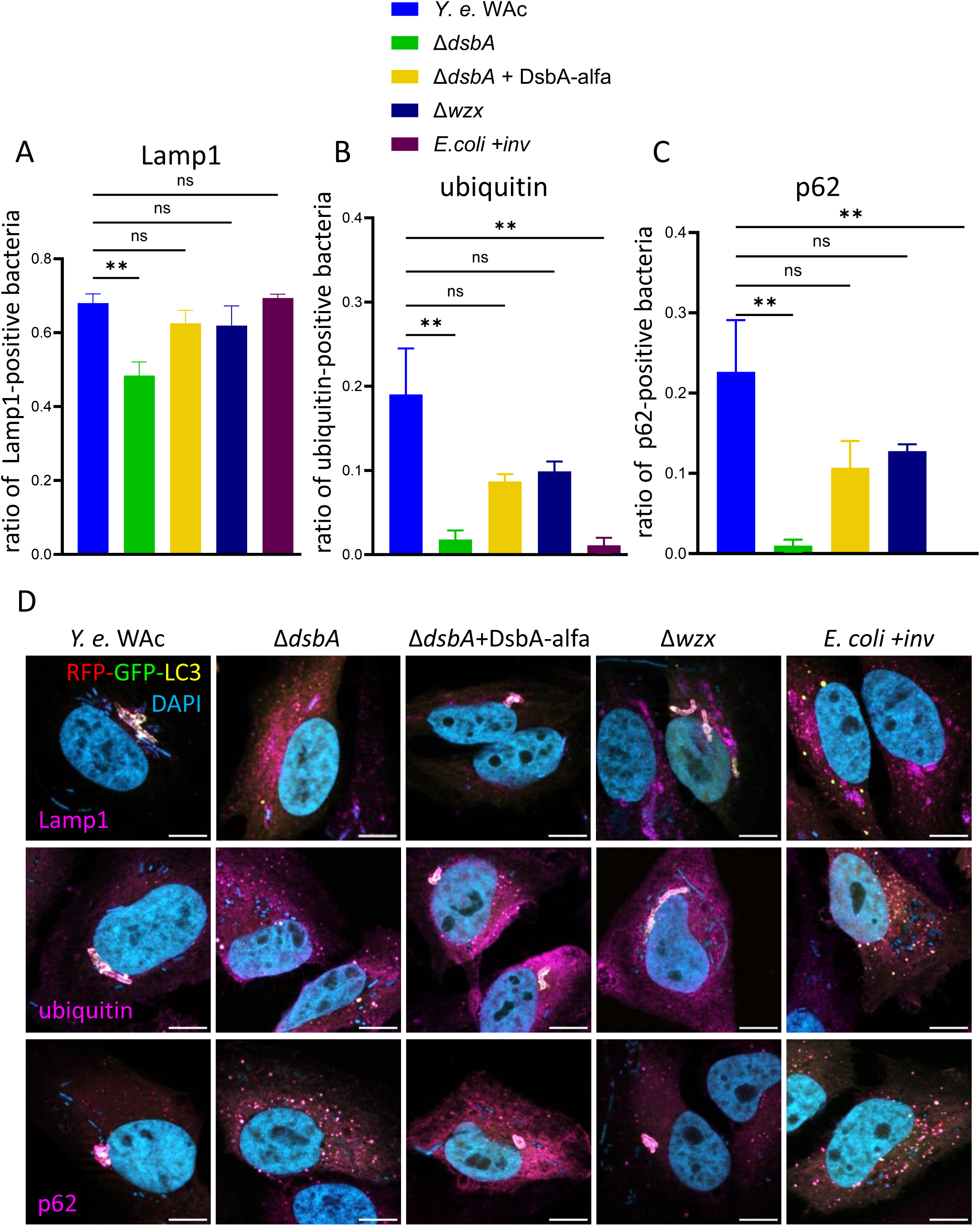
Recruitment of Lamp1 and autophagy-related adapter proteins to YCVs A. Quantification of Lamp1 positive bacteria inside HLC3 cells: HLC3-HeLa cells stably expressing RFP-GFP-LC3 cells were infected and processed for Lamp1 and DAPI staining 4 h p. i.. The numbers of DAPI-stained bacteria and bacteria inside Lamp1-positive vacuoles were counted. The ratios were calculated by dividing the numbers of Lamp1-positive bacteria by the total bacterial numbers. Three individual experiments were performed and ten images were quantified in A – C. B. Quantification of ubiquitin-positive bacteria inside HLC3 cells: HLC3 cells were infected and processed for Ubiquitin and DAPI staining as in A. C. Quantification of p62-positive bacteria inside HLC3 cells: HLC3 cells were infected and processed for p62 and DAPI staining as in A. D. Representative microscopy images of Lamp1-, ubiquitin- and p62-positive YCVs 4 h p. i.: HLC3 were infected and immunostaining of endogenous Lamp1, ubiquitin or p62 was performed. Coloured images of all channels are merged. Scale bars denote 10 μm.

Whether DsbA also plays role in intracellular survival of the virulence plasmid-bearing WA-314 strain was additionally investigated. WA-314 possesses the Ysc T3SS. The Ysc system is not required for intracellular survival of *Yersinia* and it is presently unclear whether it may fulfil any other intracellular functions and activities. We created a *dsbA* KO in the WA-314 background. The intracellular survival and replication rate of WA-314 (Fig. 6) was comparable to that of WAc (Fig. 2A), as expected. The knockout of *dsbA* in WA-314 (strain WA-314 Δ*dsbA*) severely impaired its ability to multiply intracellularly which was similar to the WAc Δ*dsbA* mutant (Fig. 2A). These findings suggest that DsbA is crucial for survival and proliferation of *Yersinia* inside human host cells irrespective of presence of the Ysc T3SS.

**Fig. 6:**
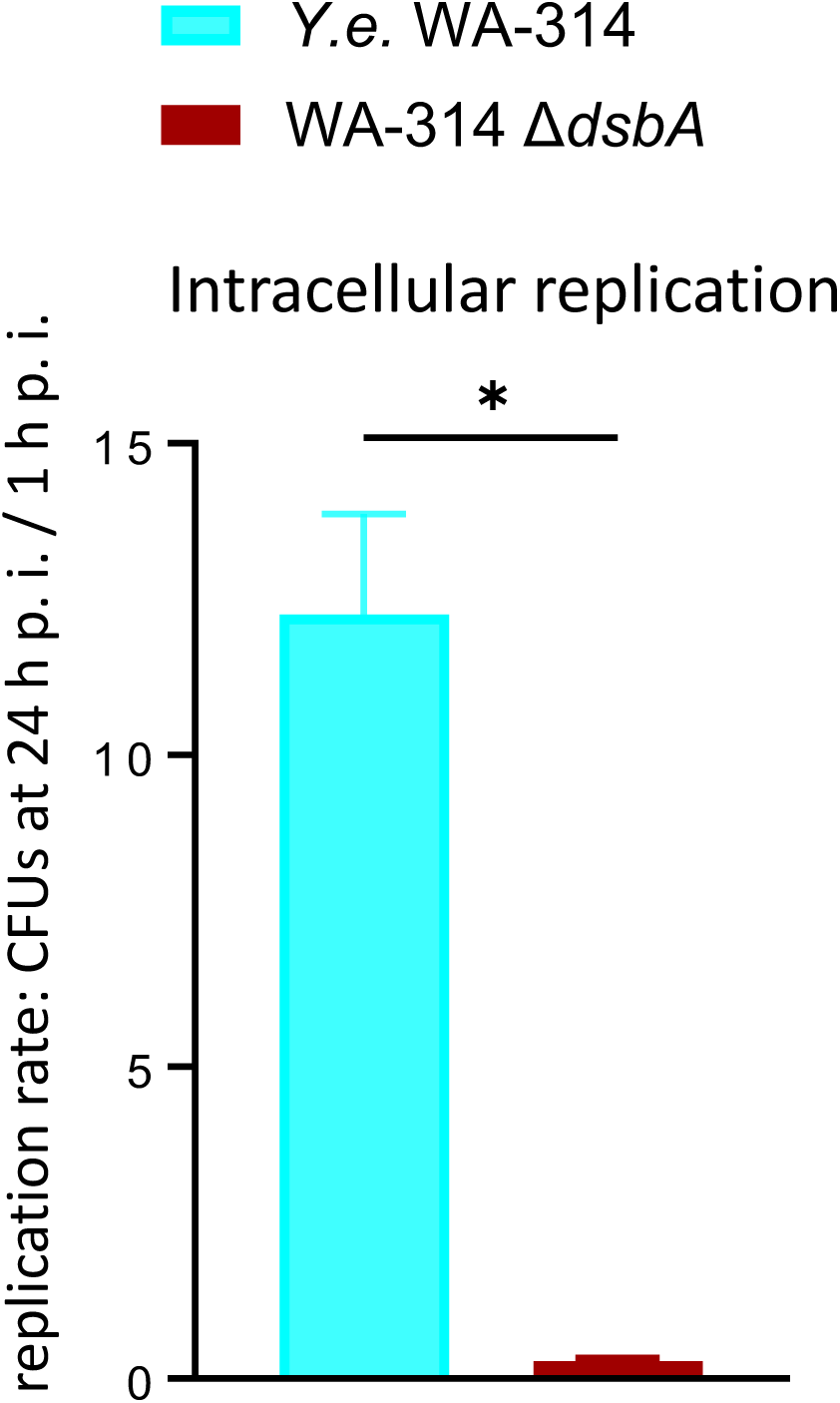
Requirement of *dsbA* for intracellular survival of pYV-harbouring wild type *Yersinia*. Analysis of intracellular survival and proliferation by gentamicin protection assay: HeLa cells were infected for 30 min, then treated with gentamicin and CFUs of intracellularly surviving bacteria were determined after 1 and 24 h p. i. The replication rates were calculated by dividing the numbers of CFUs at 24 h p. i. by the numbers of CFUs at 1 h p. i.. Three individual experiments were performed and evaluated. Welch’s T-test was performed on the data.

### AcnB plays role in inflammasome inhibition by *Yersinia* in human macrophages

A link between AcnB and inflammasome activity has previously been described for *Salmonella*. It was shown that a *Salmonella* mutant that lacks the TCA enzyme aconitase B induces rapid NLRP3 inflammasome activation in murine macrophages^30^. It is concluded that the levels of bacterial citrate, which are increased in absence of AcnB, are sensed by infected macrophages to trigger protective inflammasome activation^30^. Therefore, we wondered whether AcnB of *Yersinia* may also influence inflammasome activation in macrophages. We differentiated human monocytes to macrophages and infected them with the Δ*acnB* mutant in comparison to WAc and the other mutant strains impaired in intracellular survival. In addition, we used strain *Y.e*. pT3SS as positive control strain, which produces the *Yersinia* T3SS, but does not transfer any effector Yop^31^. This strain triggers inflammasome activation and pyroptosis through insertion of the Yop translocation pore in the macrophage membrane^32^. We found that only the Δ*acnB* mutant significantly triggered ASC speck assembly in human macrophages, which indicates inflammasome formation (Fig. 7A, 7B). This suggests that AcnB not only plays role in enabling efficient intracellular survival of *Yersinia*, but also contributes to prevent inflammasome activation. Thus, the ability of Yersinia to replicate in cells, particularly in cells that are permissive for inflammasome activation such as macrophages, may involve inhibition of inflammasome activity by AcnB.

**Fig 7:**
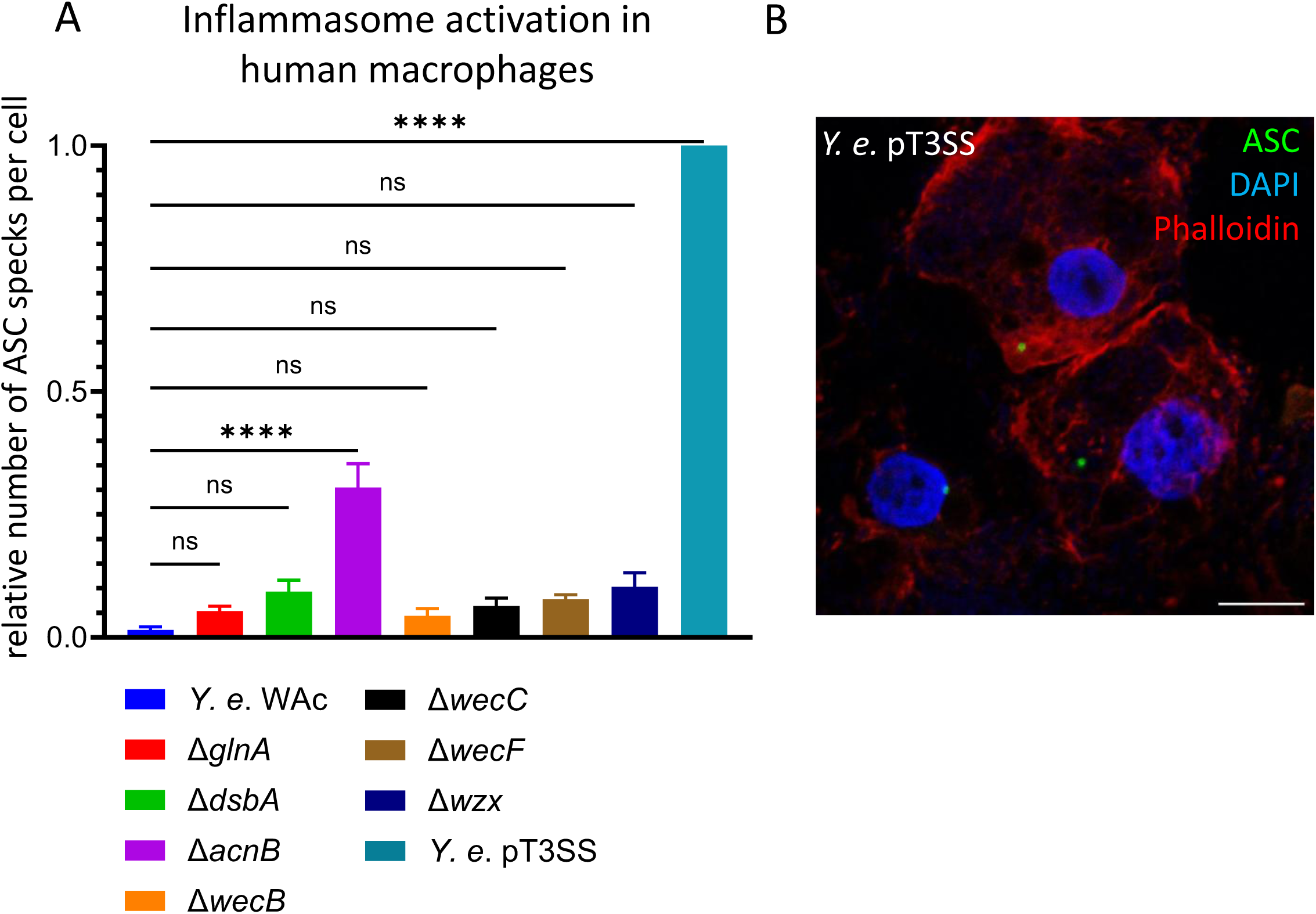
Inflammasome formation in human macrophages after *Yersinia* infection. A. Quantification of inflammasome-related ASC speck formation in human macrophages: Human monocyte-derived macrophages were infected and processed for ASC and DAPI staining 4 h p. i.. The numbers of inflammasome specks and DAPI-stained nuclei were counted. The ratios were calculated by dividing the numbers of specks by the total cell numbers. The obtained results were normalized to the ASC speck numbers induced by Y.e. pT3SS which served as positive control. Three individual experiments were performed and four images were quantified. B. Representative microscopy image for ASC speck formation: Human macrophages were infected for 4 h and immunostaining was performed to label the inflammasome adaptor protein ASC in green. Actin is stained with phalloidin in red, and DAPI marks bacterial DNA and nuclear DNA of the infected cells. Coloured images of all channels are merged. Scale bar denotes 10 μm.

## DISCUSSION

*Yersinia* is well characterized to multiply extracellularly in the lymphoid tissues of infected host organisms^1, 2^. However, there is substantial evidence that the bacteria also invade, survive, and replicate within host cells^4, 8, 9^. This is of particular importance for early infection steps, but may also contribute to persistence of the bacteria in the intestine^3, 8, 12, 33^. Because the molecular pathways and virulence factors that enable the intracellular multiplication of yersiniae are largely obscure, we applied a TraDIS approach to learn more about the virulence traits that enable the intracellular lifestyle of *Yersinia*. Host cell internalization and intracellular proliferation of *Yersinia* does not depend on the bacterial T3SS, which differs from other pathogenic gram-negative bacteria, such as *Salmonella* and *Shigella*^34^. We therefore used a *Y. enterocolitica* strain that does not have the plasmid-encoded Ysc T3SS to identify chromosomal factors involved in intracellular survival.

The obtained results indicate that it is not one particular major virulence factor that mediates the intracellular multiplication of *Yersinia* in epithelial cells. It rather appears that that ability of *Y. enterocolitica* to survive and proliferate intracellularly largely depends on several factors with pleiotropic functions in bacterial homeostasis and in adaptation to altered environmental conditions. Previous studies had already indicated that the survival of *Yersinia* inside host cells depends on bacterial stress response elements and on components of the bacterial cell envelope^4, 10, 15–21^.

In line, our findings corroborate a role of ECA structures^20^ in contributing to the intracellular proliferation of *Yersinia*. In our screen we identified three genes which encode enzymes for the biosynthesis of ECA (Table 1). These are WecB (UDP-N-acetylglucosamine 2-epimerase), WecC (UDP-N-acetyl-D-mannosamine dehydrogenase), and WecF (4-alpha-L-fucosyltransferase). The *wecB* and *wecC* genes were previously detected by Klein et al. to be important for survival of *Y. pestis* in macrophages^20^. ECA is a glycolipid produced by *Enterobacterales*. It integrates into the outer membrane leaflet along with LPS, both contributing to form the surface architecture of the bacterial envelope and a permeability barrier^35^. By that, ECA confers resistance to various external stressors and can play role in pathogenesis of infection. It has been shown that mutations or deletions in ECA synthesis genes lower the resistance of diverse *Enterobacterales* to bile salts and organic acids, reduce survival in host tissues, and attenuate pathogens in urinary tract and systemic infections^36^. Thus, ECA seems to be an important fitness factor in the host environment during infection. Our results and the findings of Klein et al.^20^ indicate that ECA also contributes to the survival of *Yersinia* in infected host cells. It is speculated that this may be due to a protective effect of ECA towards antimicrobial peptides and other harmful components in YCVs^20^.

However, besides ECA, also LPS may aid to protect against potentially toxic substances and support the intracellular replication of *Yersinia*. Our data revealed that a *Yersinia* strain defective in the LPS flippase Wzx was considerably impaired in intracellular proliferation in HeLa cells, which was similar to the *wecB* and *wecC* mutants. Wzx is a transport protein that translocates primary O-antigen repeat units across the inner bacterial membrane, then enabling assembly of the LPS coat on the bacterial surface^37^. The *wzx* knockout impairs O-antigen synthesis which may cause damage to the protective LPS barrier and enhance the sensitivity of *Y. enterocolitica* to intravacuolar toxic substances similar to a block in ECA synthesis. Also *Y. pestis* requires an intact LPS structure to be protected against host antimicrobial components and to survive intracellularly. *Y. pestis* does not produce O-antigen, but instead displays a curtailed LPS-related lipooligosaccaride structure^38^. Klein et al found that a *Y. pestis galU* mutant, which is defective in the accurate formation of lipid A and the inner core LPS region, was substantially impaired in survival in macrophages^20^. Thus, also the LPS-conferred glycolipid structure is of importance for efficient intracellular multiplication of *Yersinia*. However, it is astonishing that no other genes of the O-antigen gene cluster were identified in our TraDIS screen. The majorities of genes that direct the synthesis of LPS are organized in clusters^39^. In our transposon screen only *wzx* was identified to be of critical importance for the *Yersinia* intracellular lifestyle. This may indicate that not all genes that are conditionally essential for the intracellular survival of *Yersinia* were detected in our TraDIS assay. Or alternatively, the transposon-mediated knockout of not-identified genes required for intracellular survival of *Yersinia* was compensated by other mechanisms. Elementary vitality-preserving processes are often characterized by redundancy which may ensure adequate homeostasis-conserving cell reactions under various challenging conditions. In these cases, our TraDIS approach may have missed identification of those additionally relevant genes because their effects could have been masked or attenuated by secondary counterregulatory processes.

Interestingly, the detection of genes determining ECA and LPS synthesis as being important for the intracellular proliferation of *Yersinia* fits to a list of recently proposed core biological processes required for the full virulence of *Enterobacterales*^40^. Fouts et al. integrated genomic and Tn-Seq data to capture infection-related fitness mechanisms across multiple bacterial species. An intact ECA- and LPS-dependent carbohydrate envelope is one critical common enterobacterial fitness factor in systemic infection^40^. The bacterial polysaccharides surface structure provides protection against antimicrobial peptides, complement, and other potentially harmful host components^40^. From our data and the study of Klein et al.^20^ it appears that these suggested protective effects of ECA and LPS are also relevant for efficient intracellular survival and multiplication of *Yersinia*.

Of note, also the glutamine synthetase GlnA was proposed by Fouts et al. to belong to the infection-related core fitness factors shared by *Enterobacterales*^40^. Our TraDIS screen also revealed *glnA* to contribute to the proliferation of *Yersinia* inside host cells. GlnA converts glutamate to glutamine which are of central importance in nitrogen metabolism^41^. These amino acids are required for the synthesis of a plethora of nitrogen-containing biomolecules to ensure adequate bacterial cell growth and proliferation. Beside the importance of *glnA* in systemic enterobacterial infection, *glnA* is necessary to preserve full virulence of *Mycobacteria*^42, 43^ and *Streptococci*^44^. *GlnA* is furthermore important for the intracellular survival and replication of *Mycobacterium tuberculosis* ^43^ and *Brucella suis*^45^. It appears that these bacteria require *glnA* to thrive under nitrogen-limiting conditions inside host cells^43, 45^. *M. tuberculosis* moreover releases GlnA into the phagosome, which could influence the phagosomal pH and phagosome-lysosome fusion^46^. Such GlnA effects may also play critical roles in the intracellular lifestyle of *Yersinia*. It has been shown that the expression of *glnA* is rapidly upregulated by *Y. pestis* during replication in macrophages^47^, and that the loss of the *glnALG* operon leads to severely attenuated virulence of *Y. pestis*^48^. GlnA may thus give an impressive example for the versatile essentiality of a bacterial enzyme, being required for basic metabolism on the one hand, and for adaptation to infection and establishment of full virulence on the other.

Another enzyme that is predominantly involved in metabolic processes, but apparently also critically necessary for the adjustment of *Yersinia* to intravacuolar stress conditions during infection is aconitase B (AcnB). The knockout of *acnB* impaired productive intracellular proliferation of *Yersinia* similar to the *glnA* deficiency. AcnB is a major enzyme of the tricarboxylic acid (TCA) cycle that catalyses the reversible conversion of citrate to isocitrate^49^. It has already been found that TCA cycle-related core metabolic processes contribute to the pathogenicity of *Y. pseudotuberculosis*. *Yersinia* mutants of regulators of the TCA cycle exhibit reduced virulence, indicating importance of the pyruvate-TCA-cycle node as a control point for *Yersinia* virulence and host colonization^50^. The TCA cycle-dependent metabolism also critically governs virulence functions in *Salmonella*^51^. Furthermore, beyond its implication in the TCA cycle, AcnB may also play a role in impeding inflammasome activation. It was shown that a lack of aconitase B in *Salmonella* results in increased NLRP3 inflammasome activation in macrophages, which correlated with an attenuation of the mutant in virulence^30^. It is proposed that an increase in bacterial citrate resulting from the inactivation AcnB is sensed by infected macrophages to trigger protective inflammasome activation through mitochondrial reactive oxygen species^30^. In our experiments, the *Yersinia acnB* mutant also triggered significant inflammasome activation in human macrophages, as revealed by increased ASC speck formation. These data suggest that AcnB may specifically influence protective proinflammatory pathways besides its implication in fundamental basic and adaptive metabolic stress responses. This emphasizes AcnB as promising candidate for further exploration of its role in immune evasion and the intracellular survival of *Yersinia*.

Additional to these factors, we identified the *dsbA* gene in our TraDIS screen. *DsbA* appeared to be of major importance for the intracellular survival and multiplication of *Yersinia* in our experiments. DsbA is a periplasmic disulfide oxidoreductase which transfers disulfide bonds to unfolded proteins as they enter the periplasm^24^. This ensures the correct folding and stability of numerous extracytoplasmic, secreted and surface-exposed proteins. The substrates of DsbA also include bacterial virulence factors, such as adhesins, toxins, and secretion systems^24^. DsbA consequently controls a wide array of processes that facilitate infection and it is critically implicated in the pathogenicity of many gram-negative bacterial pathogens, i. e. of *E. coli*, *Salmonella*, *Shigella*, *Pseudomonas*, *Bordetella* and *Neisseria*^24, 52^. Also in *Yersinia*, DsbA seems to be a key regulator of virulence, as it has been shown that DsbA is required for stable expression of YscC which forms the injectisome of the *Yersinia* Ysc T3SS^53^. Loss of DsbA consequently leads to a defect in Yop secretion. The *dsbA*-negative mutant investigated in our experiments was not significantly impaired in HeLa cell adherence and invasion. However, it was most severely affected in its ability to survive and multiply in the infected cells compared to the other analyzed *Yersinia* mutants. Accordingly, *dsbA*-negative yersiniae were unable to establish intracellular infection and they were rapidly eliminated after host cell internalization. This effect occurred independent from the DsbA function in mediating proper T3SS assembly^53^, as both T3SS-positive and-negative yersiniae required *dsbA* for intracellular survival. The substrates of DsbA, which enable intracellular survival of *Yersinia*, are, however, yet unknown. Because DsbA determines the correct conformation of many proteins localized in the periplasm, inserted in the membranes, exposed to the surface, or secreted outside the bacteria, the knockout of *dsbA* may have pleiotropic effects on *Yersinia* and its virulence determinants^24, 52^. Similar as in *Pseudomonas* and *Erwinia*^54, 55^, DsbA may control several pathogenicity-associated *Yersinia* phenotypes, potentially involving surface-related protective measures and the secretion of host immunomodulatory components. Further studies are needed to characterize the DsbA-dependent *Yersinia* virulence arsenal required for intracellular survival.

Interestingly, the *dsbA* mutant was not only most significantly impaired in intracellular survival, but also in the activation of autophagic features in the infected cells. Previous studies from us and others have shown that the formation of proliferative YCVs involves the acquisition of autophagy-related characteristics^11, 13, 14^. Fusion of the autophagic YCVs with lysosomes is actively prevented by *Yersinia*^11^. It appears that yersiniae take advantage of host cell autophagy to gain access to a protected intracellular compartment for bacterial replication. For the majority of the investigated mutants our results show a close correlation between the capabilities of the strains to induce autophagy and to multiply intracellularly. Accordingly, the extent of autophagy induction by the *Yersinia* mutants positively correlated with intracellular bacterial proliferation. Deficiency for *wecB*, *wecC*, *glnA*, *acnB*, and *dsbA* affected autophagy activation as well as intracellular multiplication of the respective mutants. This indicates that these factors are critically involved in enabling both these presumably connected events in Yersinia infection. The *dsbA* knockout displays the most pronounced phenotype at this juncture. Only the *wzx* negative mutant differed in this apparent coherence, as it induced a strong autophagic response, but was nevertheless efficiently eliminated by the infected cells. Noteworthy, the *wzx* mutant-containing vacuoles were characterized by enhanced acidification in the course of infection. We therefore speculate that the *wzx* mutant is inefficient in preventing fusion of autophagic YCVs with lysosomes, which could then result in bacterial killing. This may point out a role of *wzx* and Wzx-regulated LPS synthesis processes in preventing autophagosome-lysosome fusion in *Yersinia* infection. However, also this idea requires further experimental validation.

In summary, our study suggests that the induction of autophagy and the intracellular survival and replication of *Yersinia* result from a multifactorial adaptation process of the bacteria to the intravacuolar environment. This involves the accurate formation of a protective carbohydrate surface envelope, the metabolic activities of several enzymes required for fundamental bacterial homeostasis, and likely additional, yet to be identified proteins folded by DsbA. The detailed interplay between these different *Yersinia* components and their concerted influence on the host cell will be specified in future studies.

## MATERIALS AND METHODS

### Bacterial strains, cell culture and infection conditions

The base for the *Yersinia* strains generated and investigated in this study were the *Y. enterocolitica* serogroup O8 strains WA-314 and WAc, which is the virulence plasmid-cured derivative of WA-314^56^. *E. coli* strain HB101-pINV1914 expressing the *Yersinia inv* gene (*E. coli* +*inv*)^25^ was used as control. Strain *Y. e.* T3SS harbors the segment of the pYV virulence plasmid that encodes the secretion / translocation apparatus of WA-314, but no Yop effector gene^31^. For cell infection, bacteria were grown overnight to the stationary phase in Luria-Bertani medium at 27°C (*Y. enterocolitica*) or at 37°C (*E. coli*), and bacteria were adjusted to the desired OD with DPBS.

HeLa cells were cultured at 37°C and 5% CO_2_ in Dulbeccós Modified Eagle Medium (DMEM) (Gibco™ / Thermo Fisher Scientific) supplemented with 1% penicillin/streptomycin (Thermo Fisher Scientific) and 10% fetal bovine serum (FBS) (Gibco™ / Thermo Fisher Scientific). HeLa cells stably expressing an RFP-GFP-LC3 construct (HeLa-Difluo™ hLC3 Cells HLC3 cells, InvivoGen) were cultivated in a medium containing additionally 100 μg/ml of zeocin and 100 μg/ml normocin.

Human peripheral blood monocytes were freshly isolated from buffy coats as described by Linder et al.^57^ and cultivated in RPMI medium 1640 (1X) (Gibco™ / Thermo Fisher Scientific) supplemented with 1% penicillin/streptomycin and 20% autologous serum at 37°C and 5% CO_2_. The cells were differentiated into macrophages until day 7 after isolation and then used for infection.

For infection, the cells were incubated in antibiotic free medium. HeLa cells were infected with bacteria at a MOI of 100 for gentamicin protection assays, a MOI of 200 for western blots, and a MOI of 50 (staining of intra- and extracellular bacteria and autophagy-related proteins) or 100 (lysotracker staining) for microscopy. If applicable, to synchronize infection, the bacteria were centrifuged onto the cells. 30 min later, 100 µg/ml gentamicin was added to kill the extracellular bacteria. Human monocyte-derived macrophages were infected at a MOI of 300 and gentamicin was omitted in the subsequent incubation period.

### Analysis of intracellular bacterial replication and fluorescence microscopy

Survival and replication of *Y. enterocolitica* and *E. coli* +*inv* within infected cells was analysed by gentamicin protection assay. At the given time points after infection, the cells were washed three times with DPBS and then lysed with 0.2 % Triton X-100. Intracellular bacteria were released during incubation on ice for 5 min. Serial dilutions of the bacterial suspensions were plated on agar dishes to determine the numbers of surviving intracellular bacteria. Dishes were incubated for 24 h and the numbers of colony-forming units (CFUs) were counted in three different dilutions per strain and time point.

For microscopy, a 12 mm diameter coverslip was placed in the wells prior to cell seeding. Lysotracker staining of infected cells was performed by adding Lysotracker Red dye (Invitrogen) at a final concentration of 50 nM to the cells 30 min prior to fixation at 24 h p. i.. The samples were fixed after two washes with DPBS by a 10 min incubation in 4 % PFA. If applicable, immunofluorescence staining was performed. To label extracellular bacteria, they were stained with anti-*Yersinia*-LPS antibody prior to permeabilization without the use of Triton X-100. The cell membranes were permeabilised with 0.1 % Triton X-100 for 15 min. The primary antibodies (anti-*Y. enterocolitica* O:8, Sifin Diagnostics GmbH; anti-ubiquitinylated proteins, clone FK2, Merck Millipore; p62/SQSTM1 P0067, Merck Millipore; Lamp1 D2D11, Cell Signaling Technology; ASC (B-3): sc-514414, Santa Cruz Biotechnology) were diluted in 3 % BSA with 0.05 % triton in DPBS. The secondary antibodies chicken anti-rabbit IgG AlexaFluor™ 488 (Invitrogen), donkey anti-mouse IgG AlexaFluor™ 488 (Bio-Techne GmbH), or goat anti-rabbit IgG STAR-RED 647 (Abberior GmbH) were applied together with phalloidin AlexaFluor 647 (Invitrogen), if needed. After washing, the coverslips were fixed on slides using a mounting medium containing DAPI (Invitrogen / Thermo Fisher Scientific). The Zeiss Apotome fluorescence microscope was used with a 20x objective or a 63x oil immersion objective and the ZEN 3.4 (Zen Pro) software. ImageJ was then utilized to analyse the fluorescence microscopy images. The fluorescence signals of the LPS-and Lysotracker-stained bacteria were measured to calculate the percentages of acidified bacteria by dividing the Lysotracker-positive fluorescence signals by the total LPS-positive signals. A similar procedure was applied to determine the proportion of extracellular bacteria in relation to the total population of cell-associated bacteria in double immunofluorescence staining. To analyse recruitment of host cell proteins to YCVs, the numbers of DAPI-stained bacteria and of LC3-, ubiquitin-, p62- or Lamp1-positive vacuoles, as well as the numbers of bacteria within them, were counted. The ratio of labelled bacteria was determined by dividing the LC3-, ubiquitin-, p62- or Lamp1-positive bacteria by the total number of DAPI-stained bacteria.

### SDS-PAGE and Western Blotting

At a specific time point after infection, the cells were harvested and lysed by resuspension in 4x Laemmli Sample Buffer (BioRad) and denatured at 95 °C for 10 min. Protein separation was performed by SDS-PAGE with Mini PROTEAN® Tetra Cell system (BioRad). The membrane was blocked with 5% (w/v) milk powder in TBST (15 mM Tris-HCl, 140 mM NaCl, 0.1 % (v/v) Tween20®) and incubated in a solution of the specific primary antibody LC3B 2775S rabbit (Cell Signaling Technology) and actin mouse 8H10D10 (Cell Signaling Technology) or HRP conjugated alfa-tag-antibody (NanoTag Biotechnologies) in 5 % BSA in TBST. The secondary antibodies Goat α mouse IgG (H+L) HRP conjugated (Jackson Immuno Research Laboratories Inc.) and rabbit IgG HRP linked Antibody #7074 (Cell Signaling Technology) were applied in 5 % milk in TBST. FemtoLUCENT PLUS-HRP Kit (G-Biosciences) was used to detect the target proteins with the imager ImageQuant 800, Amersham Bioscience / Cytiva. Fiji ImageJ software was used to analyse band patterns. The intensities of cytosolic LC3B-I (16 kDa) and autophagy-related lipidated LC3B-II (14 kDa) were measured and the LC3 conversion ratios were calculated by dividing the LC3B-II values by those for LC3B-I.

### Disc antimicrobial susceptibility test

A bacterial colony was resuspended in PBS and adjusted to a McFarland turbidity of 0.5 (∼1.5 × 10⁸ bacteria/ml). The suspension was evenly plated on Mueller Hinton agar. An antibiotic disc containing 10 µg gentamicin (Mast Diagnostica) was placed in the middle of each agar plate. After overnight incubation at 37 °C, the zone of inhibition diameters around the disc, reflecting the sensitivity of the investigated strain towards gentamicin, was measured.

### TraDIS mutagenesis and infection experiment

For transposon mutagenesis, the EZ-Tn5 <KAN-2> transposon (EZ-Tn5™ <KAN 2>Tnp Transposome™ Kit, Biosearch Technologies, LGC, Lucigen Epicentre) was amplified, subcloned and electroporated into *Yersinia* to allow DNA methylation. The transposon-containing plasmid was isolated, the transposon cut from the plasmid using PvuII restriction enzyme and purified. The methylated, strain-adapted transposon was then combined with the transposase enzyme, following the manufacturer’s protocol (Biozym Scientific), creating a stable transposome complex. The transposome complex was electroporated into the required WAc *Yersinia* strain so that the transposon could integrate into the genome and randomly mutate the bacterium. The resulting mutant colonies were harvested, and resuspended in 10% glycerol. The combined library contained approximately 100,000 mutants. Whole genome sequencing in a subpopulation of 20 clones showed that these mutants had only one transposon sequence integrated into the genome. The library was directly used as the first sample for sequencing (initial library). For the second sample, the bacteria were grown in liquid culture for 2 h (pre-infection library). The mutant library was then used to infect HeLa cells at an MOI of 20 and gentamicin was added to kill extracellular bacteria after one hour of infection. In parallel, cell culture medium (without eukaryotic cells) was inoculated with the bacteria as control sample and the culture was grown for 24 h (sample 3: outgrowth control library). After a final infection time of 24 hours, infected HeLa cells were washed and scraped, and then lysed with 10 ml of 0.2 % Triton X-100 to release the intracellular bacteria. The viable intracellular bacteria were regrown in culture for 3 h at 37 °C. They were then harvested and used for sequencing of the intracellular mutant library (sample 4).

### Library preparation and sequencing

Genomic DNA of the bacteria was isolated using MagNA Pure system. The samples were vortexed and stored on ice for 10–15 min. DNA shearing was performed with a Bioruptor (NGS Facility of the Leibniz Institute of Virology, Hamburg) by sonication at 4°C, consisting of six cycles of 5 s sonication with a 90 s pause between each cycle. Further library preparation was done according to the NEBNext® Ultra™ II DNA Library Prep Kit for Illumina protocol (New England BioLabs). The adapter-ligated DNA was purified using AMPure XP beads (Beckman Coulter GmbH) at a bead-to-sample ratio of 0.155x. Enrichment PCR with 22 cycles was performed to amplify transposon-containing fragments, as detailed in the kit manual, but an equal mixture of four, transposon-specific, primers (tcgtcggcagcgtcAGATGTG-TATAAGAGACAGNNNGCATGCAAGCTTCAGGGTTGA, tcgtcggcagcgtcAGATGTGTATAAGAGACAGNNNc-GCATGCAAGCTTCAGGGTTGA, tcgtcggcagcgtcAGATGTGTATAAGAGACAGNNNtaGCATGCAAGCTTCAG-GGTTGA, tcgtcggcagcgtcAGATGTGTATAAGAGACAGNNNattGCATGCAAGCTTCAGGGTTGA) was used as the forward primer. The fragments were purified again using AMPure XP beads at a ratio of 0.9x and 50 ng of the samples were used for a further index PCR of four cycles, ligating specific i7 and i5 primers for amplicon sequencing. After a final purification, the libraries were measured on TapeStation (Agilent Technologies). Amplicon sequencing was performed at the Leibniz Institute of Virology using NextSeq2000 (Illumina) sequencer and the XLEAP 300 Cycle Kit. Paired-end reads were sequenced in 300 cycles at a depth of 10 million reads per sample.

### CRISPR Cas12a-assisted, targeted mutagenesis of *Yersinia*

The CRISPR Cas12a protocol was adapted to generate *Y. enterocolitica* knockout mutants^58^. A guide RNA containing BsaI overhangs (fw: TAGAT and rev: AGAC) and a recognition sequence for the CRISPR Cas enzyme (5’-TTN-3’) was cloned into the backbone of the pAC-crRNA plasmid. It was transformed into *E. coli* Top10 and purified. The crRNA plasmid and a homology fragment were electroporated into the *Yersinia* strain carrying the CPF1-Cas12a plasmid. The homology DNA fragments were 100 bp ssDNA oligonucleotides containing a combination of two 50 bp fragments of the genomic sequences located upstream and downstream of the target gene for knockout. The CPF1 and pAC-crRNA plasmids were cured from the resulting mutants by heat stress (40°C) and by using plates containing 5% sucrose, respectively.

### Data Analysis

Raw paired-end sequencing data in FASTQ format were processed using the Transposon Position Profiling (TPP) pipeline, adapted for Tn5 transposon specificity. The analysis was performed using the reference genome of *Yersinia enterocolitica* subsp. *enterocolitica* WA-314 (GenBank accession: CP009367). Read 1 (R1) was screened for the Tn5-specific primer sequence (AGCTTCAGGGTTGAGATGTGTATAAGAGACAG), allowing up to one nucleotide mismatch. Genomic DNA flanking the transposon insertion site was extracted from R1 and R2 reads, and paired-end reads were aligned to the reference genome using BWA-MEM. Only properly paired reads mapping to opposite strands were retained for further analysis. Unique insertion events were quantified after collapsing PCR duplicates. Reads were grouped according to barcode sequence and mapping coordinates, and each unique combination was counted as a single template. Template counts at each genomic position were exported in .wig format for downstream statistical analysis. Statistical analysis of insertion patterns was conducted using Transit. Datasets were normalized using the Trimmed Total Reads (TTR) method to correct for differences in library complexity and sequencing depth. Conditional essentiality was assessed by analysis of variance (ANOVA), followed by Benjamini-Hochberg false discovery rate (FDR) correction (α = 0.05), comparing insertion counts across four experimental conditions: initial mutant library, pre-infection pool, outgrowth control, intracellular mutants. Gene-length normalization was performed by calculating the insertion density per potential insertion site to prevent longer genes from appearing artificially essential. Positional effects were taken into account by distinguishing between internal and terminal gaps in insertion coverage.

Constitutive essentiality was evaluated independently using the Tn5Gaps algorithm, which identifies genes containing significant runs of non-insertions by permutation testing. Genome-wide insertion distributions and essential gene locations were visualized using Circos. Scatter plots represented normalized template counts for each condition, and an inner heatmap highlighted genes classified as essential (FDR-adjusted *p*-value < 0.05, Tn5Gaps). All analyses were performed on the complete reference genome CP009367 to ensure accurate coordinate mapping.

### Statistics

If not indicated otherwise, statistical analyses were performed in GraphPad Prism using a one-way analysis of variance (ANOVA) with a p-value below 0.05 considered as statistically significant. Mean and standard errors of the mean (SEM) are displayed in graphics. ns= p > 0.05; *= p ≤ 0.05; **= p ≤ 0.01; ***= p ≤ 0.001; ****= p ≤ 0.0001).

## ACKNOWLEDGEMENTS

This study was funded by the Deutsche Forschungsgemeinschaft (DFG, German Research Foundation) GRK2771 – project no. 453548970 “Humans and Microbes: Reorganisation of Cell Compartments and Molecular Complexes during Infection”. We thank Joscha List for providing micrographs for Fig. 2C which he had prepared during his bachelor thesis at our institution. We are also grateful to the UKE Microscopy Imaging Facility (UMIF) for help with microscopy and image analysis. Furthermore, we thank Dr. Frank Bentzien and the Institute for Transfusion Medicine at the UKE for supply with buffy coats required for monocyte preparation, the NGS facility of the Leibniz Institute of Virology for helpful instructions, and members of the Aepfelbacher group for macrophage preparation and for providing antibodies and expertise on inflammasomes. Fig. 1A was created using BioRender.

## DATA AVAILABILITY

The next-generation sequencing (NGS) data generated in this study, including raw FASTQ files and processed insertion-site counts, have been deposited in the Gene Expression Omnibus (GEO) under the Series accession number GSE348789 (Sample accessions GSM10080565–GSM10080568; SRA accessions SRX35435089–SRX35435092).

