## Supplemental Data for "Genes controlling polysaccharide synthesis, protein folding and basic metabolism ensure intra-host cell adaptation and replication of *Yersinia enterocolitica*"

SUPPLEMENTARY FIGURE LEGENDS

**Supplementary Fig. 1: Growth properties of *Yersinia* mutants in culture media and assessment of gentamicin sensitivity.**

A. Growth curves at 27°C in LB medium and at 37°C in cell culture (DMEM) medium: The media were inoculated and the OD600 was measured hourly from 1 h to 8 h, and at 24 h. The final ODs after 24 h of growth at 37°C in DMEM medium were statistically analysed (lower panel). Three individual experiments were performed.

B. Disc antimicrobial susceptibility testing for gentamicin: The sensitivities of the *Yersinia* WT strain WAc and the WAc knockout mutants towards gentamicin were evaluated by measuring the zone of inhibition diameters around the gentamicin disk. The susceptibilities of all investigated strains were similar, with inhibition zone diameters of approximately 25 mm.

**Supplementary Fig. 2: Rates of cell association and invasion of *Yersinia* mutants.**

A. Rates of cell association: The CFU values of the HeLa cell gentamicin protection assays obtained after 1 h p. i. were used to assess initial cell association and invasion by the mutants.

B. Invasion success rates: Double immunofluorescence staining of infected HeLa cells was performed after 1 h p. i. to differentially label extracellular bacteria (green) before membrane permeabilization, and the total pool of all cell-associated bacteria (red). The fluorescence signals of the bacteria stained with the anti-LPS antibody in the different colours were measured. Successful internalization of the bacteria by the HeLa cells is expressed by a comparable low ratio of extracellular bacteria which was calculated by dividing the fluorescence results of extracellular bacteria by the fluorescence of the total bacterial population. Three individual experiments were performed and five images were quantified.

C. Representative microscopy images for cell invasion: HeLa were infected for 1 h and immunostaining with anti-LPS antibody was performed to label extra- and all cell associated bacteria in the different colours as in B. Actin is stained with phalloidin in red, and DAPI marks bacterial DNA and nuclear DNA of the infected cells. Coloured images of all channels are merged. Scale bars denote 10 µm.

Suppl. Fig. 1

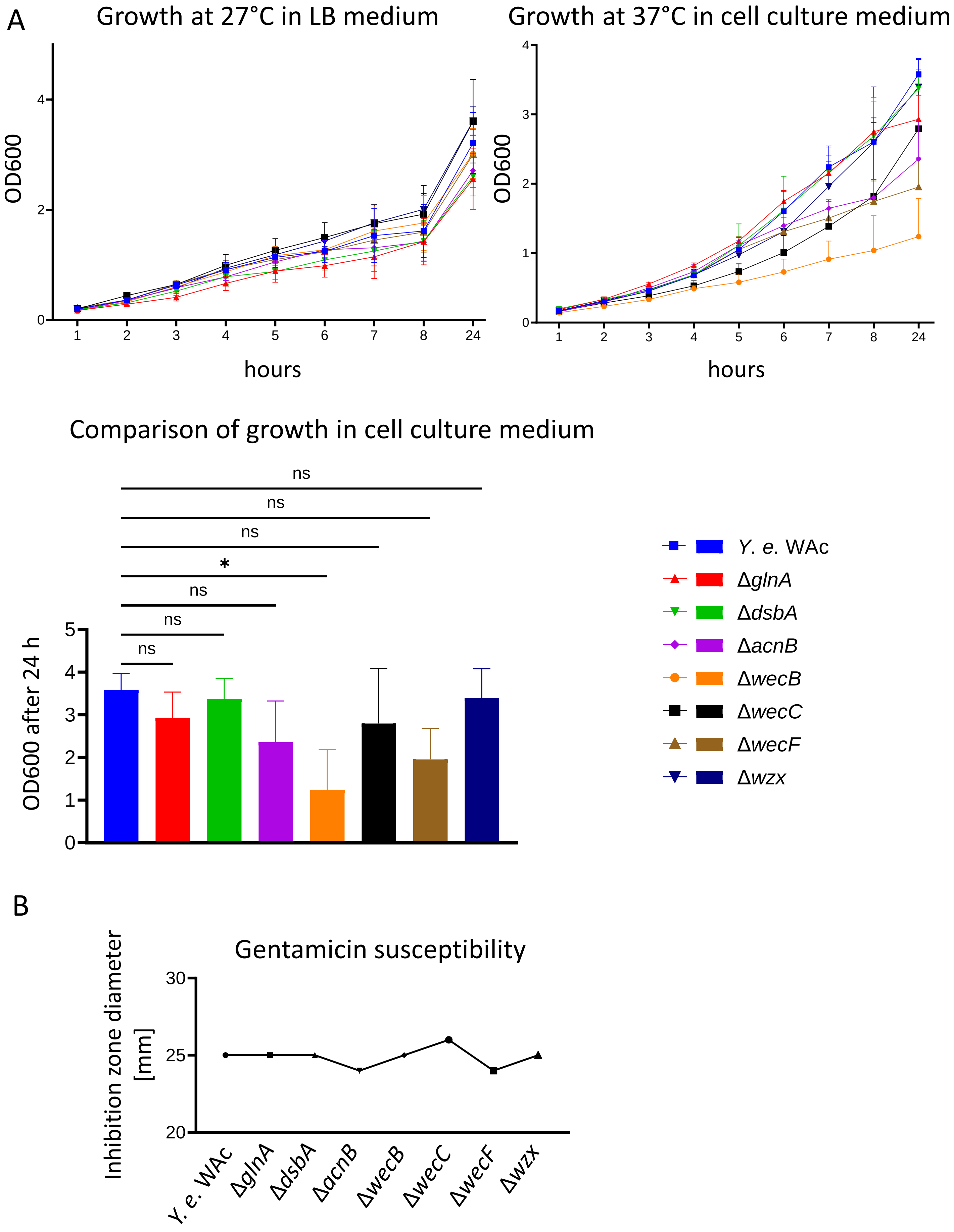
A
Growth at 27°C in LB medium
OD600
hours
Growth at 37°C in cell culture medium
OD600
hours
Comparison of growth in cell culture medium
ns
ns
ns
*
ns
ns
ns
OD600 after 24 h
Y. e. WAc
ΔglnA
ΔdsbA
ΔacnB
ΔwecB
ΔwecC
ΔwecF
Δwzx
B
Gentamicin susceptibility
Inhibition zone diameter [mm]
Y. e. WAc
ΔglnA
ΔdsbA
ΔacnB
ΔwecB
ΔwecC
ΔwecF
Δwzx

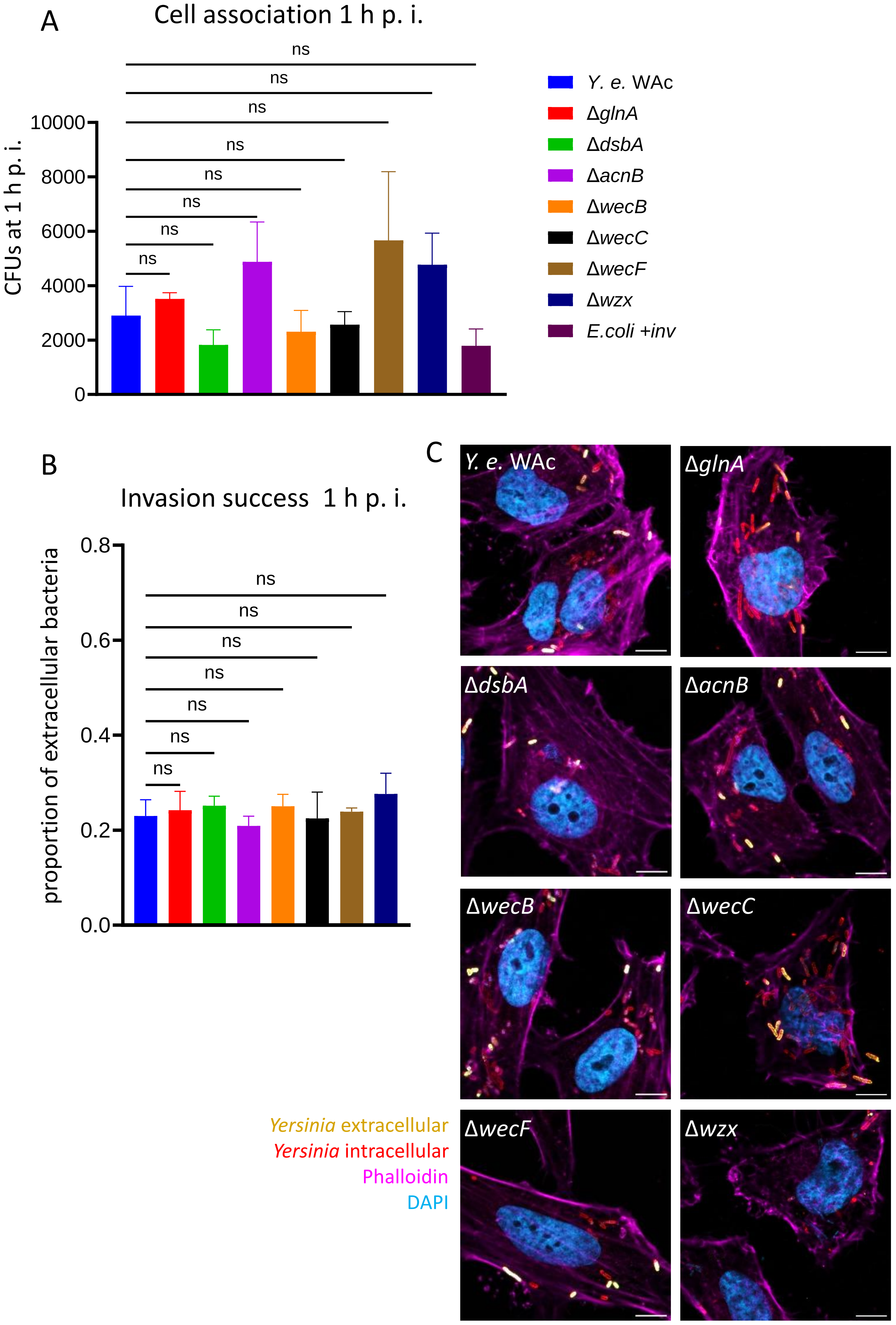

A
Cell association 1 h p. i.
ns
CFUs at 1 h p. i.
10000
8000
6000
4000
2000
0
Y. e. WAc
ΔglnA
ΔdsbA
ΔacnB
ΔwecB
ΔwecC
ΔwecF
Δwzx
E.coli +inv
B
Invasion success 1 h p. i.
proportion of extracellular bacteria
0.8
0.6
0.4
0.2
0.0
C
Yersinia extracellular
Yersinia intracellular
Phalloidin
DAPI

### SUPPLEMENTARY TABLES

**Supplementary table 1: Overview of the high-throughput sequencing results**

The number of individual insertion sites, the total number of sequenced reads and the proportion of transposon-containing sequences were evaluated per sample

| Sample No. | Sample name | Number of sequenced reads | Number of transposon-containing reads | Percentage of transposon-containing reads | Number of insertion sites |
|---|---|---|---|---|---|
| 1 | Initial library | 49,821,406 | 20,339,623 | 40.8% | 85,950 |
| 2 | Pre-infection library | 43,486,192 | 22,631,019 | 52.0% | 85,216 |
| 3 | Outgrowth control library | 70,663,823 | 26,777,280 | 37.9% | 82,414 |
| 4 | Intracellular mutant library | 51,244,639 | 23,204,461 | 45.3% | 70,705 |

**Supplementary table 2: List of the identified constitutively essential genes of *Y. enterocolitica.***

A total of 217 genes were identified as not mutated in the initial mutant library of the TraDIS screen, demonstrating the necessity of these genes for the overall survival of *Yersinia*. Raw p-values from the ANOVA F-test were corrected using the Benjamini-FDR method. The resulting adjusted p-value was used to determine essentiality (p-value <0.05). Gene-length normalization was performed and positional effects were taken into account. Constitutive essentiality was assessed by identifying genes containing significant non-insertion segments using permutation tests.

| ORF | Name | Description | COG Letters | Functional Groups | p-value adjusted |
|---|---|---|---|---|---|
| CH47_1002 | *mukF* | *mukF* | Cell cycle control | Cellular Processes | 0 |
| CH47_1003 | *mukE* | *mukE* | Cell cycle control | Cellular Processes | 0 |
| CH47_1004 | *mukB* | *mukB* | Cell cycle control | Cellular Processes | 0 |
| CH47_1011 | *asnS* | *asnS* | Translation, ribosomal structure | Information Storage | 0 |
| CH47_1076 | - | *rne* | Translation, ribosomal structure | Information Storage | 0 |
| CH47_1083 | *fabD* | *fabD* | Lipid metabolism | Metabolism | 0.00089 |
| CH47_1084 | *fabG* | *fabG* | Lipid metabolism, Secondary metabolites | Metabolism | 0 |
| CH47_1085 | *acpP* | *acpP* | Lipid metabolism, Secondary metabolites | Metabolism | 0 |
| CH47_1090 | *holB* | *holB* | Replication, recombination, repair | Information Storage | 0.00013 |
| CH47_1101 | *thiK* | *thiK* | Cell wall/membrane biogenesis | Cellular Processes | 0.0494 |
| CH47_1104 | *ndh* | *ndh* | Energy production | Metabolism | 0.03062 |
| CH47_1107 | *lolC* | *lolC* | Cell wall/membrane biogenesis | Cellular Processes | 0.00002 |
| CH47_1108 | *lolD* | *lolD* | Defense mechanisms | Cellular Processes | 0 |
| CH47_1109 | *lolE* | *lolE* | Cell wall/membrane biogenesis | Cellular Processes | 0 |
| CH47_1119 | *trmU* | *mnmA* | Translation, ribosomal structure | Information Storage | 0 |
| CH47_1207 | - | - | Function unknown | Poorly Characterized | 0.02045 |
| CH47_1217 | - | - | Unassigned | Unassigned | 0.02045 |
| CH47_1218 | *dinJ* | *dinJ* | Replication, recombination, repair | Information Storage | 0.02045 |

| | | | | | |
|---|---|---|---|---|---|
| CH47_1261 | *pgsA* | *pgsA* | Lipid metabolism | Metabolism | 0.00013 |
| CH47_1300 | *yebR* | *yebR* | Signal transduction | Cellular Processes | 0.03956 |
| CH47_1310 | *ogl* | *ogl* | Intracellular trafficking | Cellular Processes | 0.00007 |
| CH47_1314 | *ydjM* | *ydjM* | Function unknown | Poorly Characterized | 0.00148 |
| CH47_132 | *metK* | *metK* | Coenzyme metabolism | Metabolism | 0.00002 |
| CH47_1330 | - | *scpA* | Lipid metabolism | Metabolism | 0.01884 |
| CH47_1346 | *thrS* | *thrS* | Translation, ribosomal structure | Information Storage | 0 |
| CH47_1347 | *infC* | *infC* | Translation, ribosomal structure | Information Storage | 0 |
| CH47_1348 | *rpmI* | *rpmI* | Translation, ribosomal structure | Information Storage | 0.00011 |
| CH47_1349 | *rplT* | *rplT* | Translation, ribosomal structure | Information Storage | 0.00011 |
| CH47_1350 | *pheS* | *pheS* | Translation, ribosomal structure | Information Storage | 0.0036 |
| CH47_1351 | *pheT* | *pheT* | Translation, ribosomal structure | Information Storage | 0 |
| CH47_1352 | *ihfA* | *himA* | Transcription | Information Storage | 0 |
| CH47_1402 | *acnA* | *acnA* | Energy production | Metabolism | 0.01512 |
| CH47_1410 | *pyrF* | *pyrF* | Nucleotide metabolism | Metabolism | 0.03956 |
| CH47_1436 | *apxIB* | *apxIB_3* | Defense mechanisms | Cellular Processes | 0.0128 |
| CH47_144 | *pgk* | *pgk* | Nucleotide metabolism | Metabolism | 0 |
| CH47_145 | *fbaA* | *fbaA* | Carbohydrate metabolism | Metabolism | 0.00665 |
| CH47_1476 | - | - | Function unknown | Poorly Characterized | 0.00035 |
| CH47_1493 | - | *srfB* | Function unknown | Poorly Characterized | 0 |
| CH47_1502 | *rbsK* | *rbsK_1* | Coenzyme metabolism | Metabolism | 0.0001 |
| CH47_1509 | *fnr* | *fnr* | Transcription | Information Storage | 0.00883 |
| CH47_1525 | - | *azoR* | Lipid metabolism | Metabolism | 0.02242 |
| CH47_1562 | - | - | - | - | 0.0494 |
| CH47_157 | - | *visC* | Energy production, Coenzyme metabolism | Metabolism | 0.03785 |

| | | | | | |
|---|---|---|---|---|---|
| CH47_1571 | *tyrS* | *tyrS* | Translation, ribosomal structure | Information Storage | 0 |
| CH47_1578 | - | *ydhJ* | Defense mechanisms | Cellular Processes | 0.00565 |
| CH47_1579 | - | *ydhK* | Function unknown | Poorly Characterized | 0.0351 |
| CH47_1610 | - | *ydiJ* | Energy production | Metabolism | 0.03184 |
| CH47_1633 | *topA* | *topA* | Replication, recombination, repair | Information Storage | 0.00006 |
| CH47_1667 | *adhE* | *adhE* | Energy production | Metabolism | 0 |
| CH47_1688 | *ansA* | *ansA* | Amino acid metabolism, Translation, ribosomal structure | Information Storage; Metabolism | 0.00565 |
| CH47_1692 | *gap* | *gapA* | Carbohydrate metabolism | Metabolism | 0 |
| CH47_1695 | - | *yeaD* | Carbohydrate metabolism | Metabolism | 0.01402 |
| CH47_1730 | *yeaZ* | *yeaZ* | Posttranslational modification | Cellular Processes | 0 |
| CH47_175 | *lysS* | *lysS* | Translation, ribosomal structure | Information Storage | 0 |
| CH47_1751 | *aspS* | *aspS* | Translation, ribosomal structure | Information Storage | 0 |
| CH47_1755 | *cmoA* | *cmoA* | Secondary metabolites | Metabolism | 0.00103 |
| CH47_1756 | *cmoB* | *cmoB* | Translation, ribosomal structure | Information Storage | 0.00103 |
| CH47_1759 | *argS* | *argS* | Translation, ribosomal structure | Information Storage | 0 |
| CH47_1762 | *mviN* | *murJ* | Intracellular trafficking | Cellular Processes | 0.00018 |
| CH47_1778 | *kdsA* | *kdsA* | Cell wall/membrane biogenesis | Cellular Processes | 0.00001 |
| CH47_1783 | *hemA* | *hemA* | Coenzyme metabolism | Metabolism | 0.00006 |
| CH47_1784 | *lolB* | *lolB* | Cell wall/membrane biogenesis | Cellular Processes | 0.01057 |
| CH47_1785 | *ispE* | *ispE* | Nucleotide metabolism | Metabolism | 0.00007 |

| | | | | | |
|---|---|---|---|---|---|
| CH47_1788 | *pth* | *pth* | Translation, ribosomal structure | Information Storage | 0.00027 |
| CH47_1897 | - | *flgH* | Cell motility | Cellular Processes | 0.00015 |
| CH47_1898 | *flgG* | *flgG* | Cell motility | Cellular Processes | 0.00015 |
| CH47_1931 | - | - | - | - | 0.00003 |
| CH47_1932 | *cspE* | *cspC* | Transcription | Information Storage | 0.00003 |
| CH47_1965 | - | *irp3* | Secondary metabolites | Metabolism | 0.01884 |
| CH47_2067 | - | *cbiG* | Coenzyme metabolism | Metabolism | 0.00073 |
| CH47_2102 | - | *lsrB_2* | Carbohydrate metabolism | Metabolism | 0.00012 |
| CH47_2136 | *metG* | *metG* | Translation, ribosomal structure | Information Storage | 0 |
| CH47_2150 | *hyfB* | *hyfB* | Energy production, Inorganic ion metabolism | Metabolism | 0.00652 |
| CH47_2161 | - | *fdsA* | Energy production | Metabolism | 0.00062 |
| CH47_2291 | *tolB* | *tolB* | Intracellular trafficking | Cellular Processes | 0.0209 |
| CH47_2334 | - | *fldA* | Energy production | Metabolism | 0.00009 |
| CH47_2339 | *glnS* | *glnS* | Translation, ribosomal structure | Information Storage | 0 |
| CH47_2357 | *lnt* | *lnt* | Cell wall/membrane biogenesis | Cellular Processes | 0.00467 |
| CH47_2364 | *leuS* | *leuS* | Translation, ribosomal structure | Information Storage | 0 |
| CH47_2366 | *holA* | *holA* | Replication, recombination, repair | Information Storage | 0.00001 |
| CH47_2426 | *lpxH* | *lpxH* | Function unknown | Poorly Characterized | 0.01336 |
| CH47_2463 | *hemH* | *hemH* | Coenzyme metabolism | Metabolism | 0.01308 |
| CH47_2464 | *adk* | *adk* | Nucleotide metabolism | Metabolism | 0.00035 |
| CH47_2468 | *dnaX* | *dnaX* | Coenzyme metabolism | Metabolism | 0 |
| CH47_2526 | *xseB* | *xseB* | Replication, recombination, repair | Information Storage | 0.00176 |
| CH47_2527 | *ispA* | *ispA* | Coenzyme metabolism | Metabolism | 0.00176 |
| CH47_2528 | *dxs* | *dxs* | Nucleotide metabolism | Metabolism | 0 |

| | | | | | |
|---|---|---|---|---|---|
| CH47_2533 | *ribD* | *ribD* | Coenzyme metabolism | Metabolism | 0 |
| CH47_2535 | *secF* | *secF* | Intracellular trafficking | Cellular Processes | 0.00007 |
| CH47_2536 | *secD* | *secD* | Intracellular trafficking | Cellular Processes | 0.00002 |
| CH47_259 | *rpsB* | *rpsB* | Translation, ribosomal structure | Information Storage | 0.00003 |
| CH47_260 | *tsf* | *tsf* | Translation, ribosomal structure | Information Storage | 0.01253 |
| CH47_263 | *dxr* | *dxr* | Lipid metabolism | Metabolism | 0 |
| CH47_2641 | *rimM* | *rimM* | Translation, ribosomal structure | Information Storage | 0.01138 |
| CH47_2655 | *alaS* | *alaS* | Translation, ribosomal structure | Information Storage | 0 |
| CH47_266 | *rseP* | *rseP* | Cell wall/membrane biogenesis | Cellular Processes | 0.00025 |
| CH47_267 | *yaeT* | *bamA* | Cell wall/membrane biogenesis | Cellular Processes | 0 |
| CH47_271 | *lpxA* | *lpxA* | Lipid metabolism | Metabolism | 0.00005 |
| CH47_2719 | *ispD* | *ispD* | Lipid metabolism | Metabolism | 0.00062 |
| CH47_272 | *lpxB* | *lpxB* | Lipid metabolism | Metabolism | 0.0313 |
| CH47_2738 | *eno* | *eno* | Carbohydrate metabolism | Metabolism | 0 |
| CH47_274 | *dnaE* | *dnaE* | Replication, recombination, repair | Information Storage | 0 |
| CH47_2771 | *can* | *can* | Inorganic ion metabolism | Metabolism | 0.00354 |
| CH47_278 | *tilS* | *tilS* | Translation, ribosomal structure | Information Storage | 0.00041 |
| CH47_2800 | *secA* | *secA* | Intracellular trafficking | Cellular Processes | 0.00005 |
| CH47_2803 | *lpxC* | *lpxC* | Cell wall/membrane biogenesis | Cellular Processes | 0.0108 |
| CH47_2804 | *ftsZ* | *ftsZ* | Cell cycle control | Cellular Processes | 0.00001 |
| CH47_2805 | *ftsA* | *ftsA* | Cell cycle control | Cellular Processes | 0.02194 |
| CH47_2806 | *ftsQ* | *ftsQ* | Cell cycle control | Cellular Processes | 0 |
| CH47_2808 | *murC* | *murC* | Cell wall/membrane biogenesis | Cellular Processes | 0.00001 |
| CH47_2810 | *ftsW* | *ftsW* | Cell cycle control | Cellular Processes | 0.01103 |

| | | | | | |
|---|---|---|---|---|---|
| CH47_2811 | *murD* | *murD* | Cell wall/membrane biogenesis | Cellular Processes | 0.00111 |
| CH47_2812 | *mraY* | *mraY* | Cell wall/membrane biogenesis | Cellular Processes | 0.00981 |
| CH47_2813 | *murF* | *murF* | Cell wall/membrane biogenesis | Cellular Processes | 0 |
| CH47_2814 | *murE* | *murE* | Cell wall/membrane biogenesis | Cellular Processes | 0 |
| CH47_2815 | *ftsI* | *ftsI* | Cell wall/membrane biogenesis | Cellular Processes | 0 |
| CH47_2849 | *lptD* | *lptD* | Cell wall/membrane biogenesis | Cellular Processes | 0.00001 |
| CH47_285 | *proS* | *proS* | Translation, ribosomal structure | Information Storage | 0.00514 |
| CH47_2862 | *ispH* | *ispH* | Lipid metabolism, Cell wall/membrane biogenesis | Cellular Processes; Metabolism | 0.02418 |
| CH47_2866 | *ribF* | *ribF* | Coenzyme metabolism | Metabolism | 0.00012 |
| CH47_2871 | *dnaK* | *dnaK* | Posttranslational modification | Cellular Processes | 0 |
| CH47_2976 | *lptG* | *lptG* | Function unknown | Poorly Characterized | 0.00013 |
| CH47_2977 | *lptF* | *lptF* | Function unknown | Poorly Characterized | 0.03184 |
| CH47_2980 | *valS* | *valS* | Translation, ribosomal structure | Information Storage | 0 |
| CH47_3041 | *rimP* | *rimP* | Translation, ribosomal structure | Information Storage | 0.00503 |
| CH47_3045 | *glmM* | *glmM* | Carbohydrate metabolism | Metabolism | 0 |
| CH47_3047 | *ftsH* | *ftsH* | Posttranslational modification | Cellular Processes | 0 |
| CH47_3054 | *cgtA* | *obg* | Function unknown | Poorly Characterized | 0 |
| CH47_3055 | - | *yhbE* | Amino acid metabolism, Carbohydrate metabolism | Metabolism | 0 |
| CH47_3058 | *ispB* | *ispB* | Coenzyme metabolism | Metabolism | 0.00589 |

| | | | | | |
|---|---|---|---|---|---|
| CH47_3081 | *rpsR* | *rpsR* | Translation, ribosomal structure | Information Storage | 0.01163 |
| CH47_3120 | *groL* | *groL* | Posttranslational modification | Cellular Processes | 0 |
| CH47_3121 | *groS* | *groS* | Posttranslational modification | Cellular Processes | 0.00012 |
| CH47_3190 | *rpoC* | *rpoC* | Transcription | Information Storage | 0 |
| CH47_3191 | *rpoB* | *rpoB* | Transcription | Information Storage | 0 |
| CH47_3192 | *rplL* | *rplL* | Translation, ribosomal structure | Information Storage | 0 |
| CH47_3193 | *rplJ* | *rplJ* | Translation, ribosomal structure | Information Storage | 0 |
| CH47_3195 | *rplK* | *rplK* | Translation, ribosomal structure | Information Storage | 0.00006 |
| CH47_3206 | *murB* | *murB* | Cell wall/membrane biogenesis | Cellular Processes | 0 |
| CH47_3227 | *ubiB* | *ubiB* | Coenzyme metabolism | Metabolism | 0.01968 |
| CH47_3257 | *rpoH* | *rpoH* | Transcription | Information Storage | 0 |
| CH47_3260 | *ftsY* | *ftsY* | Cell cycle control | Cellular Processes | 0 |
| CH47_3317 | *rho* | *rho* | Nucleotide metabolism | Metabolism | 0.00004 |
| CH47_3434 | *coaBC* | *coaBC* | Coenzyme metabolism | Metabolism | 0.01711 |
| CH47_3463 | *spoT* | *spoT* | Transcription, Signal transduction | Cellular Processes; Information Storage | 0 |
| CH47_3483 | *hemN* | *hemN* | Coenzyme metabolism | Metabolism | 0 |
| CH47_3550 | *dnaA* | *dnaA* | Replication, recombination, repair | Information Storage | 0 |
| CH47_3551 | *dnaN* | *dnaN* | Replication, recombination, repair | Information Storage | 0 |
| CH47_3553 | *gyrB* | *gyrB* | Replication, recombination, repair | Information Storage | 0 |
| CH47_3558 | *dcm* | *dcm* | Replication, recombination, repair | Information Storage | 0 |
| CH47_3577 | *glyQ* | *glyQ* | Translation, ribosomal structure | Information Storage | 0 |

| | | | | | |
|---|---|---|---|---|---|
| CH47_3578 | *glyS* | *glyS* | Translation, ribosomal structure | Information Storage | 0.0001 |
| CH47_3763 | *trpS* | *trpS* | Translation, ribosomal structure | Information Storage | 0 |
| CH47_3801 | *rpsL* | *rpsL* | Translation, ribosomal structure | Information Storage | 0 |
| CH47_3802 | *rpsG* | *rpsG* | Translation, ribosomal structure | Information Storage | 0 |
| CH47_3803 | *fusA* | *fusA* | Translation, ribosomal structure | Information Storage | 0.00842 |
| CH47_3812 | *rplB* | *rplB* | Translation, ribosomal structure | Information Storage | 0 |
| CH47_3813 | *rpsS* | *rpsS* | Translation, ribosomal structure | Information Storage | 0 |
| CH47_3814 | *rplV* | *rplV* | Translation, ribosomal structure | Information Storage | 0 |
| CH47_3815 | *rpsC* | *rpsC* | Translation, ribosomal structure | Information Storage | 0 |
| CH47_3816 | *rplP* | *rplP* | Translation, ribosomal structure | Information Storage | 0 |
| CH47_3817 | *rpmC* | *rpmC* | Translation, ribosomal structure | Information Storage | 0 |
| CH47_3819 | *rplN* | *rplN* | Translation, ribosomal structure | Information Storage | 0 |
| CH47_3820 | *rplX* | *rplX* | Translation, ribosomal structure | Information Storage | 0 |
| CH47_3821 | *rplE* | *rplE* | Translation, ribosomal structure | Information Storage | 0 |
| CH47_3822 | - | *rpsN* | Translation, ribosomal structure | Information Storage | 0 |
| CH47_3823 | *rpsH* | *rpsH* | Translation, ribosomal structure | Information Storage | 0 |
| CH47_3824 | *rplF* | *rplF* | Translation, ribosomal structure | Information Storage | 0 |

| | | | | | |
|---|---|---|---|---|---|
| CH47_3825 | *rplR* | *rplR* | Translation, ribosomal structure | Information Storage | 0 |
| CH47_3829 | *secY* | *secY* | Intracellular trafficking | Cellular Processes | 0.00896 |
| CH47_3830 | *rpmJ* | *rpmJ* | Translation, ribosomal structure | Information Storage | 0.00896 |
| CH47_3831 | *rpsM* | *rpsM* | Translation, ribosomal structure | Information Storage | 0 |
| CH47_3832 | *rpsK* | *rpsK* | Translation, ribosomal structure | Information Storage | 0 |
| CH47_3833 | *rpsD* | *rpsD* | Translation, ribosomal structure | Information Storage | 0 |
| CH47_3834 | *rpoA* | *rpoA* | Transcription | Information Storage | 0 |
| CH47_3835 | *rplQ* | *rplQ* | Translation, ribosomal structure | Information Storage | 0 |
| CH47_3877 | *plsB* | *plsB* | Lipid metabolism | Metabolism | 0 |
| CH47_3948 | *dnaB* | *dnaB* | Replication, recombination, repair | Information Storage | 0 |
| CH47_4040 | *lptA* | *lptA* | Function unknown | Poorly Characterized | 0 |
| CH47_4041 | *lptC* | *lptC* | Function unknown | Poorly Characterized | 0 |
| CH47_4042 | *kdsC* | *kdsC* | Nucleotide metabolism | Metabolism | 0 |
| CH47_4051 | *murA* | *murA* | Cell wall/membrane biogenesis | Cellular Processes | 0 |
| CH47_4052 | *degS* | *degS* | Posttranslational modification | Cellular Processes | 0.00004 |
| CH47_4116 | *rpoD* | *rpoD* | Transcription | Information Storage | 0 |
| CH47_4117 | *dnaG* | *dnaG* | Replication, recombination, repair | Information Storage | 0.00034 |
| CH47_4119 | - | *tsaD* | Translation, ribosomal structure | Information Storage | 0.03343 |
| CH47_4123 | *cca* | *cca* | Nucleotide metabolism | Metabolism | 0.00981 |
| CH47_4140 | *parE* | *parE* | Replication, recombination, repair | Information Storage | 0.00062 |
| CH47_4144 | *parC* | *parC* | Replication, recombination, repair | Information Storage | 0 |

| | | | | | |
|---|---|---|---|---|---|
| CH47_431 | *lepB* | *lepB* | Intracellular trafficking | Cellular Processes | 0 |
| CH47_453 | *nadE* | *nadE* | Coenzyme metabolism | Metabolism | 0.0163 |
| CH47_471 | *iscR* | *iscR* | Transcription | Information Storage | 0 |
| CH47_472 | *iscS* | *iscS* | Amino acid metabolism | Metabolism | 0 |
| CH47_489 | *hisS* | *hisS* | Translation, ribosomal structure | Information Storage | 0 |
| CH47_544 | *upp* | *upp* | Nucleotide metabolism | Metabolism | 0.0448 |
| CH47_552 | *dapA* | *dapA* | Amino acid metabolism | Metabolism | 0.00205 |
| CH47_561 | *dapE* | *dapE* | Amino acid metabolism | Metabolism | 0 |
| CH47_571 | *napA* | *napA* | Energy production | Metabolism | 0.00589 |
| CH47_607 | *ligA* | *ligA* | Replication, recombination, repair | Information Storage | 0.00296 |
| CH47_704 | *mnmC* | *mnmC* | Translation, ribosomal structure | Information Storage | 0.01126 |
| CH47_705 | *fabB* | *fabB* | Lipid metabolism, Secondary metabolites | Metabolism | 0.00045 |
| CH47_729 | *accD* | *accD* | Lipid metabolism | Metabolism | 0.00004 |
| CH47_731 | - | *dedD* | Cell cycle control | Cellular Processes | 0.04154 |
| CH47_801 | - | *tyrP* | Intracellular trafficking | Cellular Processes | 0.0036 |
| CH47_809 | - | *nrdA* | Nucleotide metabolism | Metabolism | 0.00966 |
| CH47_811 | *gyrA* | *gyrA* | Replication, recombination, repair | Information Storage | 0 |
| CH47_864 | - | *fruB* | Carbohydrate metabolism | Metabolism | 0.01926 |
| CH47_882 | - | - | Unassigned | Unassigned | 0.01138 |
| CH47_883 | - | *efe* | Energy production | Metabolism | 0.01138 |
| CH47_969 | *cydC* | *cydC* | Defense mechanisms | Cellular Processes | 0.00205 |
| CH47_978 | *serS* | *serS* | Translation, ribosomal structure | Information Storage | 0.00087 |
| CH47_988 | *cmk* | *cmk* | Nucleotide metabolism | Metabolism | 0 |
| CH47_989 | *rpsA* | *rpsA* | Translation, ribosomal structure | Information Storage | 0 |
| CH47_992 | *msbA* | *msbA* | Defense mechanisms | Cellular Processes | 0 |

| | | | | | |
|---|---|---|---|---|---|
| CH47_993 | *lpxK* | *lpxK* | Lipid metabolism | Metabolism | 0 |
| CH47_997 | - | *ycaR* | Function unknown | Poorly Characterized | 0.0001 |
| CH47_998 | *kdsB* | *kdsB* | Cell wall/membrane biogenesis | Cellular Processes | 0.0001 |

**Supplementary table 3: List of the identified conditionally essential genes involved in intracellular survival of *Y. enterocolitica***

Genes that were significantly less frequently mutated (p-value < 0.05) in the main sample (1) than in the pre-infection pool of mutants (2) or than in the outgrowth control (3) were termed conditional essential genes. Adjusted p-values above 0.05 were considered not significant (ns = p > 0.05).

| ORF | Name | Description | COG Letters | Functional Groups | Intracel-lular vs LB p-value adjusted | Intracel-lular vs outgrowth p-value adjusted |
|---|---|---|---|---|---|---|
| CH47_1907 | *inv* | *invasin* | Function unknown | Poorly Characterized | 0.000001 | 0 |
| CH47_3480 | *glnA* | - | Amino acid transport and metabolism | Metabolism | 0.000124 | 0.000765 |
| CH47_3307 | *wecF* | *rffT* | Energy production | Metabolism | 0.00016 | 0.006147 |
| CH47_3313 | *wecC* | - | Cell wall/membrane /envelope biogenesis | Cellular Processes | 0.00016 | 0.003165 |
| CH47_3314 | *wecB* | - | Cell wall/membrane /envelope biogenesis | Cellular Processes | 0.000447 | 0.035969 |
| CH47_3487 | *dsbA* | - | Cell wall/membrane /envelope biogenesis | Cellular Processes | 0.00056 | 0.00244 |
| CH47_2457 | *wzx* | - | Function unknown | Poorly Characterized | 0.000583 | 0.001218 |
| CH47_2780 | *acnB* | - | Energy production and conversion | Metabolism | 0.001415 | 0.004934 |
| CH47_2859 | *carB* | - | Nucleotide transport and metabolism | Metabolism | 0.000002 | n.s. |
| CH47_3090 | *purA* | - | Nucleotide transport and metabolism | Metabolism | 0.000008 | n.s. |
| CH47_3749 | *igaA* | *yrfF* | Function unknown | Poorly Characterized | 0.000008 | n.s. |
| CH47_3840 | *trkA* | - | Inorganic ion transport and metabolism | Metabolism | 0.000124 | n.s. |
| CH47_3174 | *purH* | - | Nucleotide transport and metabolism | Metabolism | 0.00016 | n.s. |

| | | | | | | |
|---|---|---|---|---|---|---|
| CH47_3518 | *atpA* | - | Nucleotide transport and metabolism | Metabolism | 0.000392 | n.s. |
| CH47_3034 | *pnp* | - | Translation, ribosomal structure and biogenesis | Information Storage | 0.00056 | n.s. |
| CH47_232 | *recB* | *recB* | Replication, recombination, repair | Information Storage | 0.000603 | n.s. |
| CH47_3390 | *priA* | *priA* | Replication, recombination, repair | Information Storage | 0.001446 | n.s. |
| CH47_3760 | *dam* | - | Coenzyme transport and metabolism | Metabolism | 0.001446 | n.s. |
| CH47_3989 | *mreC* | *mreC* | Cell wall/membrane biogenesis | Cellular Processes | 0.001446 | n.s. |
| CH47_3988 | *mreB* | *mreB* | Cell cycle control | Cellular Processes | 0.00197 | n.s. |
| CH47_3316 | *wecA* | - | Cell wall/membrane /envelope biogenesis | Cellular Processes | 0.003268 | n.s. |
| CH47_3214 | *trkH* | - | Inorganic ion transport and metabolism | Metabolism | 0.015587 | n.s. |
| CH47_3520 | *atpD* | - | Nucleotide transport and metabolism | Metabolism | 0.016488 | n.s. |
| CH47_230 | *recC* | - | Replication, recombination and repair | Information Storage | 0.018522 | n.s. |
| CH47_497 | *guaA* | *guaA* | Nucleotide metabolism | Metabolism | 0.018522 | n.s. |
| CH47_3523 | *glmS* | *glmS* | Cell wall/ membrane biogenesis | Cellular Processes | 0.019281 | n.s. |
| CH47_447 | *purL* | - | Nucleotide transport and metabolism | Metabolism | 0.020747 | n.s. |
| CH47_3486 | *polA* | *polA* | Replication, recombination, repair | Information Storage | 0.041654 | n.s. |
| CH47_3175 | *purD* | - | Nucleotide transport and metabolism | Metabolism | 0.043178 | n.s. |
| CH47_3311 | *rfbA* | *rffH* | Coenzyme transport and metabolism | Metabolism | 0.043178 | n.s. |
